# Genomic constraints to drought adaptation

**DOI:** 10.1101/2021.08.07.455511

**Authors:** Collin W Ahrens, Kevin Murray, Richard A Mazanec, Scott Ferguson, Jason Bragg, Ashley Jones, David T Tissue, Margaret Byrne, Justin O Borevitz, Paul D Rymer

## Abstract

Global shifts in climate and precipitation patterns are altering the diversity and structure of forests. The ability for species to adapt is especially difficult for long lived foundation species with unknown genetic and trait diversity. We harnessed genomic, physiological, and climate data to determine adaptation constraints. We used denovo assembly and 6.5 million genomic variants with drought related traits from 432 individuals sourced from across the complete range of the foundation tree species *Corymbia calophylla*. We found genomic variants determining traits predominantly in gene regulatory regions. The ability for populations to adapt was limited by within population genetic diversity associated with traits, and epistatic interactions within traits and pleiotropic interactions among traits. Nevertheless, we could accurately predict adaptive traits using genomic and climate data to guide reforestation. Our study indicated that some populations may contain variation sufficient for the species to adapt to the effects of drought, while other populations will need increased variability from those sources.

## Background

Climate change is predicted to shift global precipitation patterns (1), resulting in trees that are ill-equipped to respond to exacerbated droughts (2). However, many studies use climate as a proxy for phenotype, but actual phenotypic variation may be much greater than the variation predicted solely by climate. Thus, we expect that solely using climate predicted intraspecific adaptive variation will overpredict mean distribution and fail to identify within-population adaptive standing genomic variation.

The increase of drought around the world will have significant effects on long lived trees requiring adaptation of physiological traits (3). One trait, δ^13^C, is a strong indicator of drought resistance (4, 5); plants with a smaller δ^13^C are more drought resistant because they fix more carbon when water is limited (4). Other functional traits are also used as indicators of plant capacity to respond to climate fluctuations (e.g. specific leaf area (SLA), normalised difference vegetation index (NDVI), leaf size, wood density, nitrogen content, and growth). These traits have been used in many ecophysiological contexts to understand adaptive and plastic responses to environmental conditions. This work is especially important in trees, given the environmental and economic value of forested ecosystems. Although there have been studies on phenology in poplar (6) and on growth in oaks and pines (7, 8), there have been few studies that identify the genetic variants that underlie control of complex traits in response to drought events for ecologically important trees. Therefore, we developed and used genomic resources to identify intraspecific genetic variation associated with drought resistance traits in a foundational Australian tree species, *Corymbia calophylla*.

## Species and Functional Traits

Evaluation of gene-trait interactions was performed for *Corymbia calophylla* due to its ecological significance in the southwestern cape floristic region in southwestern Australia which has been undergoing dramatic changes in precipitation patterns (9). We sourced seeds from 12 populations with 10 families per population, 3-4 individuals per family to capture the species geographic and climatic distribution and characterised 432 individual trees grown in a common garden to estimate genetically determined trait variation.

We focused on three uncorrelated functional traits that are known to be indicative of drought resistance (δ^13^C (10); SLA (11); NDVI (12)). Water use efficiency (measured as carbon isotope discrimination; δ^13^C) is the link between photosynthesis and evaporation (13) that indicates climatic tolerance under water limitation (14). High specific leaf area (SLA; a ratio of leaf area to mass) indicates a large leaf surface area for water loss (transpiration) and increases tree susceptibility to drought-induced mortality (15). Leaf-level (modified red-edge) normalized difference vegetation index (NDVI; see methods) is a surrogate for chlorophyll content by quantifying leaf greenness, and can be indicative of photosynthetic activity and plant stress to drought (16), where high NDVI reflects high photosynthetic activity and low plant stress. Two of the three traits (δ^13^C and NDVI) exhibited small, but significant, narrow-sense heritabilities based on quantitative genetic models (17), indicating that some of the trait variation was controlled by polygenetic mechanisms.

We controlled for random variation of the measured trait values, using best linear unbiased predictions (BLUPs), and found that the three traits varied across families and populations (Figure 1; Figure S1). Using a linear model, analyses of BLUP trait values showed that traits were significantly differentiated among populations (F_11,419_ = 88.89; *p* < 0.001), among families within populations (F_23,407_ = 53.75; *p* < 0.001), and δ^13^C was significantly described by precipitation of the driest month (F_1,429_ = 278.7; *p* < 0.001; R^2^ = 0.39), but not significant for NDVI (F_1,425_ = 0.70; *p* = 0.40) or SLA (F_1,427_ = 0.85; *p* = 0.35).

**Figure 1.**
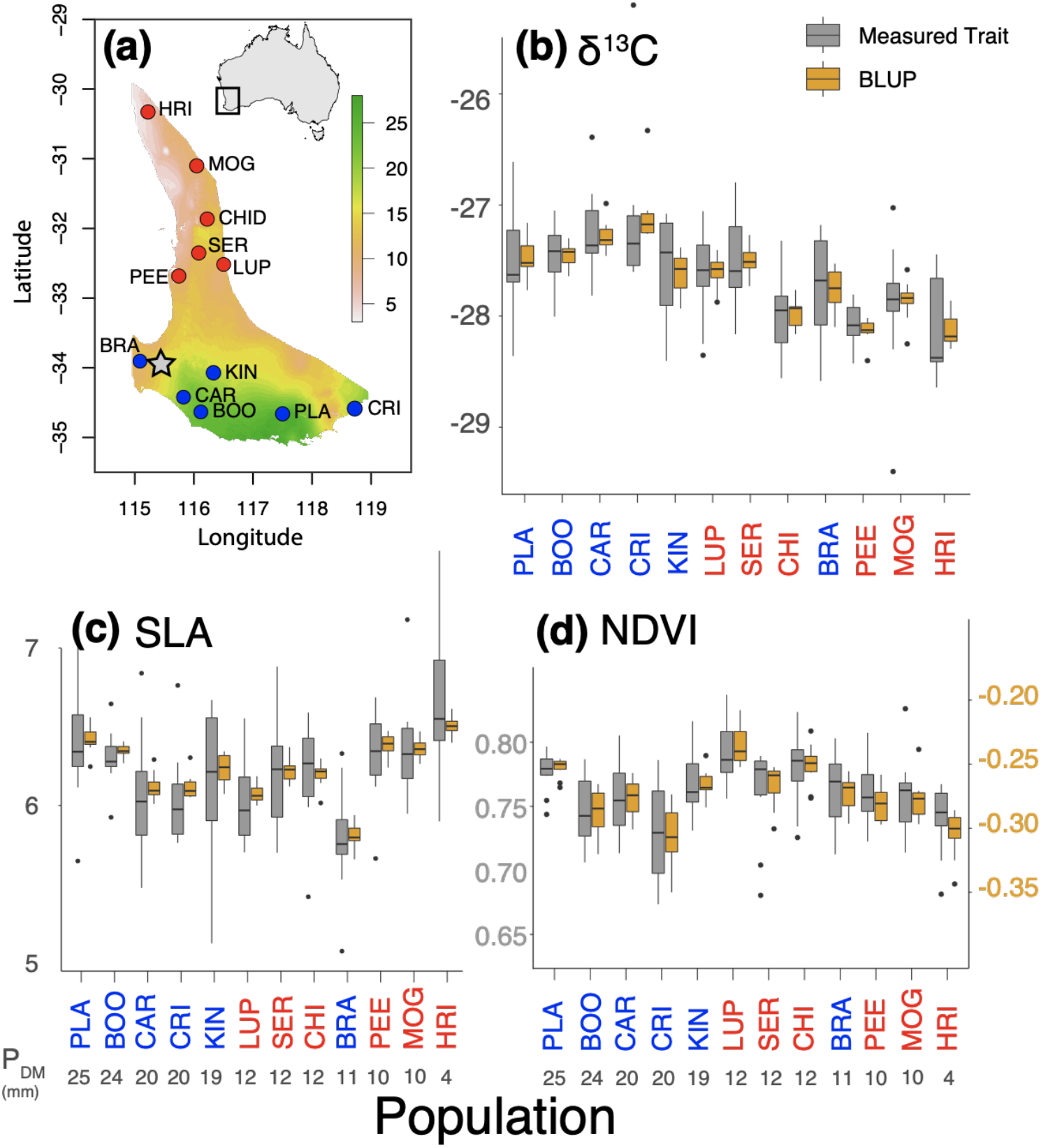
Phenotypic traits were studied in a common garden of populations sampled from across the range of *Corymbia calophylla*. (a) Population locations with precipitation of the driest month (mm; BIO14); (b-d) trait values (grey) with their best linear unbiased predictions (BLUPs; yellow) for (b) δ^13^C, (c) SLA, and (d) NDVI. Population colours are coded red for northern populations and blue for southern populations. Inset shows location of study area within Australia. Star represents the location of the experimental site. BLUP = best linear unbiased prediction. NDVI was scaled (y2-axis) to meet assumptions of normality before estimating BLUPs.

## Draft genome and genome-wide association studies (GWAS)

We extracted high molecular weight DNA from a reference *Corymbia calophylla* individual from the Australian Botanic Gardens, sequenced Blue PippinHT filtered long reads (>40kb) to 40X on two nanopore minion flow cells and assembled a denovo genome (350Mb haploid size, 100 contigs, N50 3Mb, 11 chromosome scaffolds). We then aligned individual short read sequences and identified 91M single nucleotide polymorphisms (SNPs) (Figure S2; Table S1). After filtering on quality and allele frequency, 6.49 million informative SNPs were discovered, averaging a SNP every 60 bases across all 11 chromosomes. Linkage disequilibrium (LD) decayed quickly, as median base pair distance to half-maximal *r*^2^ values were 160 base pairs (Table S2; Figure S3). These LD estimates are similar to, but slightly greater than, previous half-maximal *r*^2^ estimates of LD decay in *Eucalyptus* species (92 and 113 bp) (18), confirming that there is a very high degree of population diversity and recombination in the system.

To discover SNP-trait associations, we performed genome wide association studies (GWAS) for nine functional traits but focus on δ^13^C, NDVI, SLA (six traits not discussed: leaf size, wood density, nitrogen content, photochemical reflectance index, height, diameter; qq plots Figure S4 & manhattan plots Figure S5). There was weak but detectable population structure (*F*_ST_= 0.05). We controlled for this in the GWAS analysis using the first 10 axes of a multidimensional scaling plot (Figure S6). The resulting GWAS identified 279, 69 and 92 significant SNPs for δ^13^C, SLA, and NDVI, respectively (Figure 2a). Candidate SNPs were found on all chromosomes across the genome with several regions having a high density of candidate SNPs (e.g. Fig 2a shows peaks on 3,8,10). Magnification of the peaks in chromosomes 3, 8, and 10 show that the peaks were caused by many SNPs that occur in small regions (150-350 kb), beyond the median LD decay, interspersed with non-significant SNPs, and different among all three traits (Figure 2b). These patterns within gene rich regions could be the result of soft selective sweeps (19), where no variant sweeps to fixation because of the reliance on multiple alleles of small-effect for adaptation. Alternatively, these adaptive regions could be directly due to conserved, large haplotype blocks, which have been found to be important in adaptation among sunflower ecotypes (20). Indeed, large haplotype blocks could explain the strong LD between candidate SNPs within chromosomes and traits (R^2^ > 0.9) and within chromosomes among traits (Figure S7d-f), but not the patterns of long-range LD across different chromosomes among candidate SNPs for all three traits (Figure S7a-c). This pattern is likely due to rarity disequilibrium (a.k.a. genetic indistinguishability - giSNP (21)), a widespread phenomenon that arises when interchromosomal SNPs are in perfect LD due to the combinatorial limit on unique genotype patterns in small sample sizes. This can be alleviated with additional diversified sampling, or a backcrossing experiment to take advantage of recombination and independent assortment of giSNPs. In practice, this means that true localisation of causal variants is impossible, but at least a subset of the biological signals identified are likely real.

**Figure 2.**
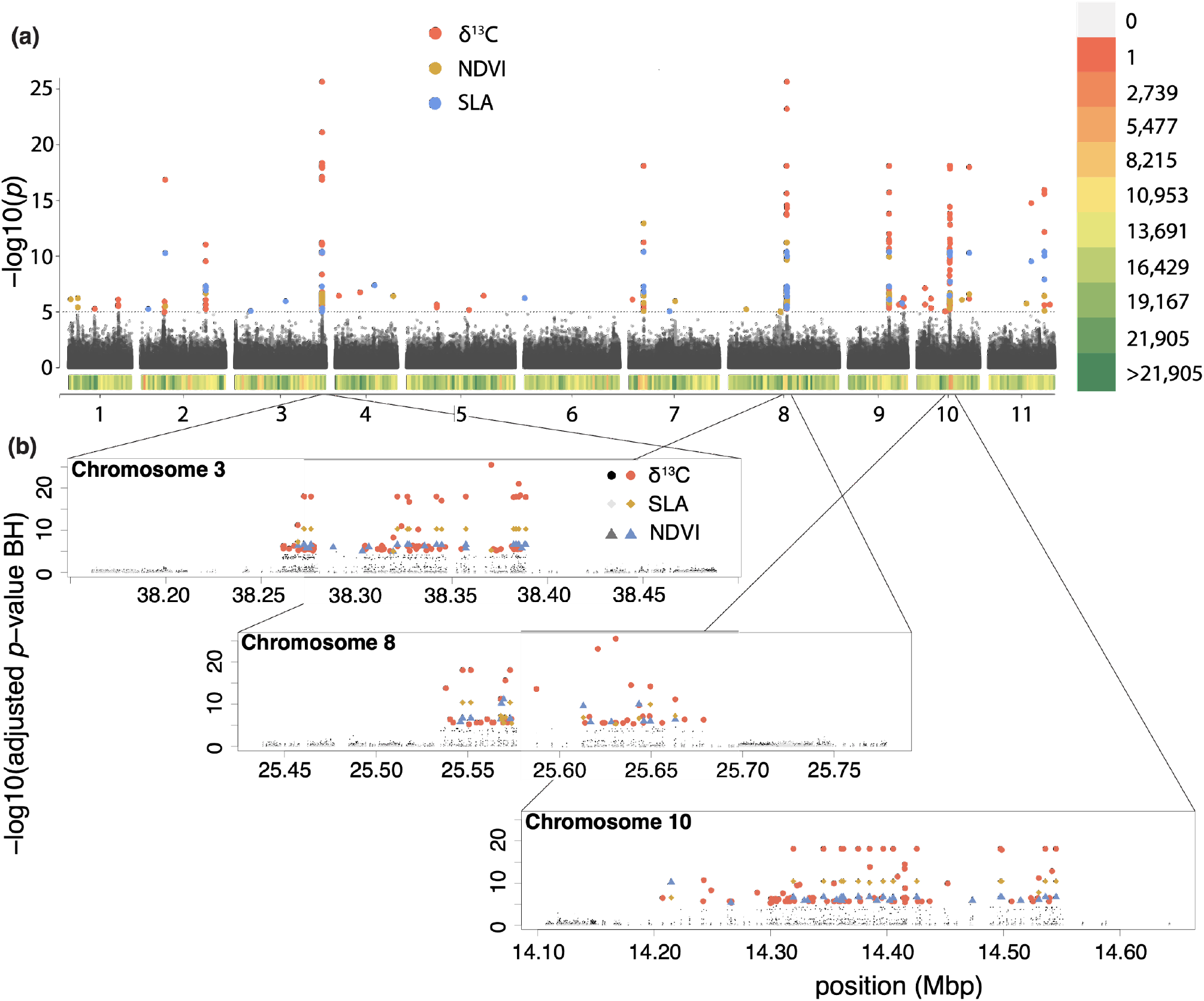
Genome sampling and GWAS outputs for *Corymbia calophylla* grown in a common environment. (a) Manhattan plots for three traits and SNP density. Inner ring is δ^13^C, the middle ring is NDVI, and the outer ring is SLA. Red points represent SNPs significantly associated with the trait at an FDR value < 0.00001. The density plots (outer coloured heatmap) are in 1 million base pair segments and colour (red to green) represents increasing density of SNPs within that segment. (b) Magnified view of the significant peaks within the three regions of adaptive variation.

To determine how much trait variation could be explained by all of the SNPs, we estimated the SNP-based heritability using common variants to explain the total proportion of variance in phenotypes (22). We found that SNP-based heritability for all three traits (**δ^13^C** *h*^2^_SNP_ = 0.55 (SE 0.14); **SLA** *h*^2^_SNP_ = 0.27 (0.12); and **NDVI** *h*^2^_SNP_ = 0.66 (0.14) was much greater than the heritabilities calculated through quantitative genetics methods (*h*^2^ = 0.11 (0.08), 0.08 (0.08), and 0.15 (0.08) for δ^13^C, SLA, and NDVI respectively; (17)). This suggests that most SNPs had relatively small effect sizes, but we were able to identify SNPs with large effects sizes. For example, top 10 candidate SNPs associated with δ^13^C showed greater effects (18 - 25%) (Table 1), than the greatest explanatory SNPs for NDVI and eight of the SNPs for SLA. There were five SNPs associated with SLA that were giSNPs, and were dropped from the combined model. Regardless, these ten SNPs accounted for ~50% of the variation for δ^13^C and SLA, and 34% of the variation in NDVI. The lower combined *R*^2^ value for NDVI compared to the other two traits might be due to many factors, including more trait variation, more SNPs of small effect, an outcome that is supported by the high SNP heritability, and more epistatic interactions.

**Table 1.** Results from independent linear models of gene trait interactions in *Corymbia calophylla* with SNP as the independent variable and phenotype as the dependent variable. The total variation explained among the top 10 SNPs in a single model. Bolded SNPs are found twice in the table. All *R*^2^ values are adjusted and significant (*p* < 0.0001). *maf* = minor allele frequency. *es* = full model effect size. VE = proportion of variation explained. ***, **, * significance in the combined model < 0.001, 0.01, 0.05 respectively. † - dropped from the combined model due to giSNP (genetically indistinguishable SNP) or too similar to another SNP.

| $\delta^{13}\text{C}_3$ | | | | SLA | | | | NDVI | | | |
| --- | --- | --- | --- | --- | --- | --- | --- | --- | --- | --- | --- |
| Variant ID | $R^2$ | *maf* | *es* | Variant ID | $R^2$ | *maf* | *es* | Variant ID | $R^2$ | *maf* | *es* |
| 10:14288468[cc] | 0.25** | 0.165 | -0.23 ± 0.03 | 8:25569614[cc] | 0.34*** | 0.497 | -0.074 ± 0.01 | 7:6588079[cc] | 0.17 | 0.416 | -0.011 ± 0.002 |
| 3:38317350[cc] | 0.24 | 0.174 | -0.19 ± 0.03 | **11:24419468[cc]** | 0.30*** | 0.135 | 0.093 ± 0.01 | 8:25645984[cc] | 0.12** | 0.494 | 0.011 ± 0.002 |
| 3:38370345[cc] | 0.21** | 0.160 | -0.22 ± 0.02 | 7:6588358[cc] | 0.09* | 0.119 | 0.074 ± 0.01 | 7:20379383[cc] | 0.11*** | 0.047 | 0.030 ± 0.004 |
| 8:25630714[cc]† | 0.21 | 0.160 | -0.23 ± 0.02 | 8:25649571[cc] | 0.08 | 0.071 | 0.10 ± 0.01 | **11:24419468[cc]** | 0.10 | 0.135 | -0.017 ± 0.002 |
| 8:25620946[cc] | 0.21 | 0.169 | -0.22 ± 0.02 | 3:38327356[cc] | 0.07 | 0.064 | 0.11 ± 0.01 | 11:16551085[cc] | 0.05 | 0.019 | -0.041 ± 0.006 |
| 8:25550760[cc] | 0.19** | 0.128 | 0.17 ± 0.03 | 3:38388540[cc] | 0.07 | 0.063 | 0.10 ± 0.01 | 1:4110088[cc] | 0.05 | 0.068 | -0.025 ± 0.003 |
| 3:38385682[cc] | 0.18* | 0.146 | -0.21 ± 0.02 | 10:14497305[cc]† | 0.07 | 0.063 | 0.10 ± 0.01 | 10:14265736[cc] | 0.05* | 0.396 | 0.013 ± 0.002 |
| 3:38332167[cc] | 0.18** | 0.166 | 0.17 ± 0.02 | 3:38272343[cc]† | 0.07 | 0.063 | 0.10 ± 0.01 | 10:14473331[cc] | 0.05 | 0.427 | 0.013 ± 0.002 |
| 10:14324927[cc] | 0.18*** | 0.455 | 0.15 ± 0.02 | 3:38275883[cc]† | 0.07 | 0.063 | 0.10 ± 0.01 | 3:38335476[cc] | 0.05 | 0.400 | 0.014 ± 0.002 |
| 9:17931039[cc] | 0.18 | 0.084 | -0.13 ± 0.02 | 3:38321179[cc]† | 0.07 | 0.063 | 0.10 ± 0.01 | 3:38306212[cc]† | 0.04 | 0.412 | 0.014 ± 0.002 |
| Combined 10 SNP $R^2$ | 0.51 | | | | 0.48 | | | | 0.34 | | |
| Full model VE | 0.55 |  |  |  | 0.27 |  |  |  | 0.66 |  |  |

Complex gene and trait interactions (i.e. epistasis and pleiotropy) are known to play important roles in quantitative traits (23). We found evidence that some of the unexplained trait variation could be attributable to epistasis based on interactions among significant SNPs identified in the GWAS. Gene interactions among the significant SNPs were explicitly evaluated using CAPE (importantly giSNPs are removed from this analysis), and there are significant epistatic interactions across the genome among the significant SNPs (*p* < 0.05; multiple testing correction using the FDR method; Figure 3a), with strong interactions between chromosomes 3, 8, 9, and 10. Main effects between two SNP pairs are shown between chromosomes 2 & 8 (negative effect; blue arrows in 3a) and 8 & 10 (positive effect; yellow arrows in 3a) (Figure 3b), and we also provide visualisation of an epistatic interaction when the main effect (variant + trait) is conditioned on a second variant (Figure 3c), where the interaction between the two SNPs effects the trait in a negative way (Figure 3c dashed line). We then assessed possible pleiotropic interactions between the three leaf-level traits, i.e. the effect of one SNP on multiple traits. Pleiotropic interactions are shown in Figure 3a in the concentric bands, where the same SNP is highlighted for more than one trait. We identified several cases within chromosomes 1, 3, 7, 8, 9, and 10 that were due to pleiotropic effects (Figures 3a & S8). While some of the interactions affected traits in the same direction, there were also antagonistic interactions present (when a gene affects traits in different directions; represented by different colours along the grey trait-circles in Figures 3 & S8). Pleiotropy was corroborated through annotation results, for example, there were 11 genes that were found to be significantly associated with all three traits. Eight of these genes were expressed during growth and development processes, including in plant structure such as guard cell and leaf structure. When assessing epistatic and pleiotropic interactions, it is difficult to surmise how single SNPs truly control functional traits. Indeed, there are 10s of thousands of pairwise possibilities in our dataset alone; we have not tried to assess the total effect of all interactions.

**Figure 3.**
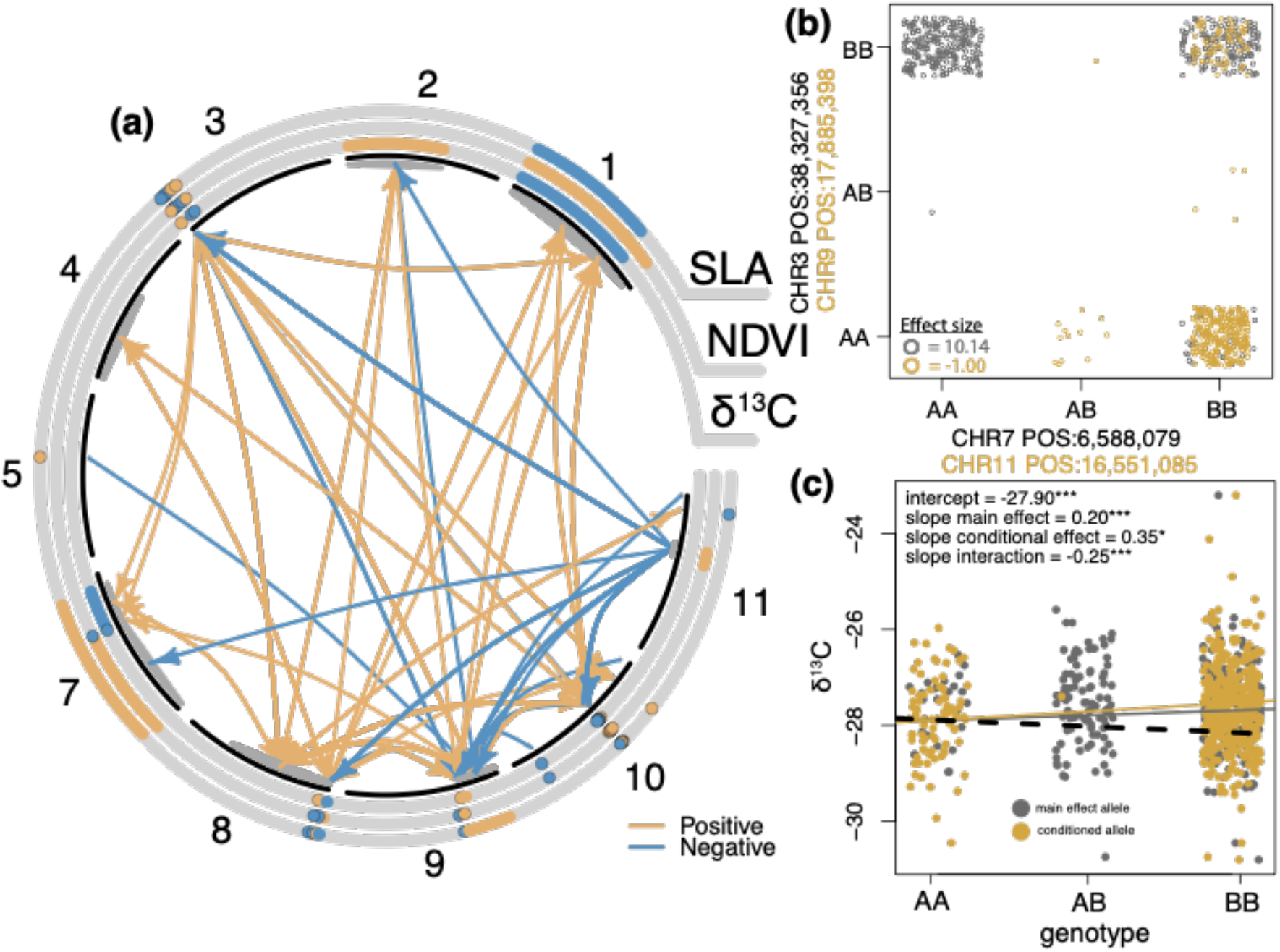
Patterns of significant gene-gene (epistasis) and gene-trait (pleiotropy) interactions. (a) Interactions between SNPs (epistasis) are shown with coloured arrows and pleiotropic effects between traits are shown in the circular bands. Chromosomes are in black, chromosome six is not present because no SNPs were significant in the analysis. The direction of influence is shown by colour, where orange indicates that the SNP affects a different SNP in a positive way and blue is indicative of a negative effect. Interactions between a SNP and multiple traits indicate pleiotropy, while the same colours are indicative of the same effects. Antagonistic pleiotropy is inferred if the colours are different among SNPs in the same chromosomal location. Points on chromosome 3, 8, and 10 have been manually separated due to severe overlapping to visualise the antagonistic effects. (b) Genotypes for negatively influenced epistatic interaction between two SNP variants (grey points in (b) & blue arrows in (a)) and positively influenced epistatic interactions between two SNP variants (orange points in (b) & orange arrows in (a)). (c) Visualisation of one significant epistatic interaction where the main effect of a SNP (grey in (c)) and trait is conditioned on a second SNP (yellow in (c)), black dashed line is the interaction effect between the two variants. *P*-value – *** < 0.001; * < 0.05.

There is also a proportion of trait variation that was not revealed. One explanation might be that these traits are highly additive, and we were only able to identify the variants most strongly associated with the phenotype; hence, many small-effect alleles related to a specific trait went undetected due to our statistically conservative approach. SNP heritability will capture these undetected small effects, but this only explained just greater than half of the variation for δ^13^C. Another explanation is that phenotypic plasticity plays a major role in these traits e.g. (24), and due to stochastic effects we could not determine some sources of variation. Indeed, plasticity is a crucial mechanism facilitating plant persistence in variable environments, and it remains a major challenge for plant ecologists and evolutionary biologists to partition trait variation into plastic or adaptive bins (25). Mechanistic properties of plastic responses are complex, but they could be due to methylation or control through regulatory elements (26). Methylation differences could result in non-heritable differences, while variants in regulatory elements could result in differences in plastic responses that are heritable. These are all areas of great interest and follow up studies could focus on these mechanisms. However, our results shown above indicate two main findings: 1) a large proportion of the variation can be explained by relatively few QTLs, and 2) traits are controlled by complex gene interactions that are difficult to quantify.

## Annotation

In order to identify location (e.g., *cis* regulatory, genic) and effect (e.g., synonymous, nonsynonymous) of adaptation, snpEFF was used to annotate all SNPs and candidate SNPs. Annotations for all 6.49 million filtered SNPs reveal many moderate (nonsynonymous) and low (synonymous) effect alleles with a much smaller proportion of high-effect alleles (Table S3). Each chromosome had similar rates of nonsynonymous and synonymous SNPs (Table S3), except for chromosome 8 with a much higher rate of SNPs upstream and downstream of genes and more than double the number of high-effect SNPs than the other chromosomes (Table S3). Candidate SNPs within the three chromosomes with the highest significant peaks show interesting patterns (Chromosomes 3, 8, and 10; red in Figure 2a). For example, of the 41 SNPs on chromosome 8 associated with δ^13^C, 37 are within gene regulatory regions (< 5,000 base pairs upstream of the gene) (Table 2), while for the four remaining SNPs, one was nonsynonymous with a moderate effect, and three were synonymous with low effects (Table 2). The regions of adaptive variation in chromosomes 3 and 10 are mostly intergenic with SNPs in gene *cis*-regulatory regions (within 5kb of a gene on either the 5’ (upstream) or 3’ (downstream) end of the gene) and six candidate SNPs in promoter regions (within 500 bp upstream of a gene) for δ^13^C on chromosome 10. This is notable because sequence variation in regulatory regions can give rise to differential regulation of nearby genes (27) and are an effective substrate for selection.

**Table 2.** Annotation summary for the significant SNPs on three chromosomes. N = number of SNPs; H = high; M = moderate; L = low effect; prom = promoter region (wihtin 500 bp of a gene); N = the number of candidate SNPs; NS = nonsynonymous; S = synonymous; Intergenic = SNP not found within 5kb of a gene.

| chromosome | trait | N | up-reg | down-reg | prom | NS | S | effect (H\|M\|L) | intergenic |
| --- | --- | --- | --- | --- | --- | --- | --- | --- | --- |
| 3 | SLA | 17 | 1 | 2 | 0 | 0 | 0 | 0\|0\|0 | 14 |
|  | NDVI | 23 | 2 | 3 | 0 | 0 | 0 | 0\|0\|0 | 18 |
| | $\delta^{13}\text{C}$ | 74 | 6 | 8 | 0 | 1 | 0 | 0\|1\|0 | 59 |
| 8 | SLA | 13 | 13 | 0 | 6 | 0 | 0 | 0\|0\|0 | 0 |
|  | NDVI | 16 | 16 | 0 | 5 | 0 | 0 | 0\|0\|0 | 0 |
| | $\delta^{13}\text{C}$ | 41 | 37 | 0 | 25 | 1 | 3 | 0\|1\|3 | 0 |
| 10 | SLA | 16 | 8 | 3 | 0 | 0 | 0 | 0\|0\|0 | 5 |
|  | NDVI | 26 | 11 | 7 | 0 | 0 | 0 | 0\|0\|0 | 8 |
| | $\delta^{13}\text{C}$ | 88 | 37 | 19 | 6 | 2 | 0 | 0\|2\|0 | 28 |

To gain a better understanding of gene functional relationships, we used GO enrichment analysis to compare pathways among traits and within processes (Figure S9). Gene networks were different among traits, in that they show different processes within the same category. For instance, the largest network in biological process for δ^13^C focuses on transport of ions, hydrogen, and nitrogen, while SLA is highlighted by defense response and NDVI is highlighted by a reduced transport network compared to δ^13^C.

## Trait predictions from genotype and climate

To investigate whether candidate SNPs and climate variables independently describe phenotypic distribution, we used random forest models separately for SNPs and climate. As input into the random forest model, we summarised each suite of candidate SNPs for the three traits separately using discriminant analysis of principal coordinates (DAPC; Figure S10). The results for δ^13^C are shown here, and NDVI and SLA are given in supplementary information (Figures S11 & S12). NDVI shows a similar pattern to δ^13^C. However, SLA is quite different and seems to lack biological relevance. This is probably due to excessive within-population variation, which is not surprising because it was found to be not heritable (17) and is known to be highly plastic (28). For δ^13^C, the first two DAPC axes explained 48 and 22% of the genomic variation (Figure S10a). Together, the two axes showed peaks and valleys in a 3d surface-plot that do not align with linear relationships between genetic variation and δ^13^C (Figure 4a). The random forest model with the two most significant climate variables (maximum temperature of the warmest month (importance = 6.68)) and precipitation of the driest month (importance = 18.22)) – from a full random forest model with six different climate factors – explains 69% of the variation and appears to have a similar pattern compared to the genomic prediction (53% of the variation; Figure 4a & S11) with one caveat; the z-axis from the climate prediction had greater breadth compared to the genomic prediction, which is more evident when plotted on the 2D map (Figure 4b). Indeed, the difference between the two predictive maps is greatest at the northern and southern regions, with an absolute difference of 0.3, which is substantial considering the trait variance across the distribution (0.5) (Figure 4b). These differences can be attributed to heritable variation, where the genomic variants only account for heritable variation while the climate factors attempt to explain all trait variation, including variation due to plasticity and dominance effects for example. This indicates that climate predictions likely overpredicted adapted variation.

**Figure 4.**
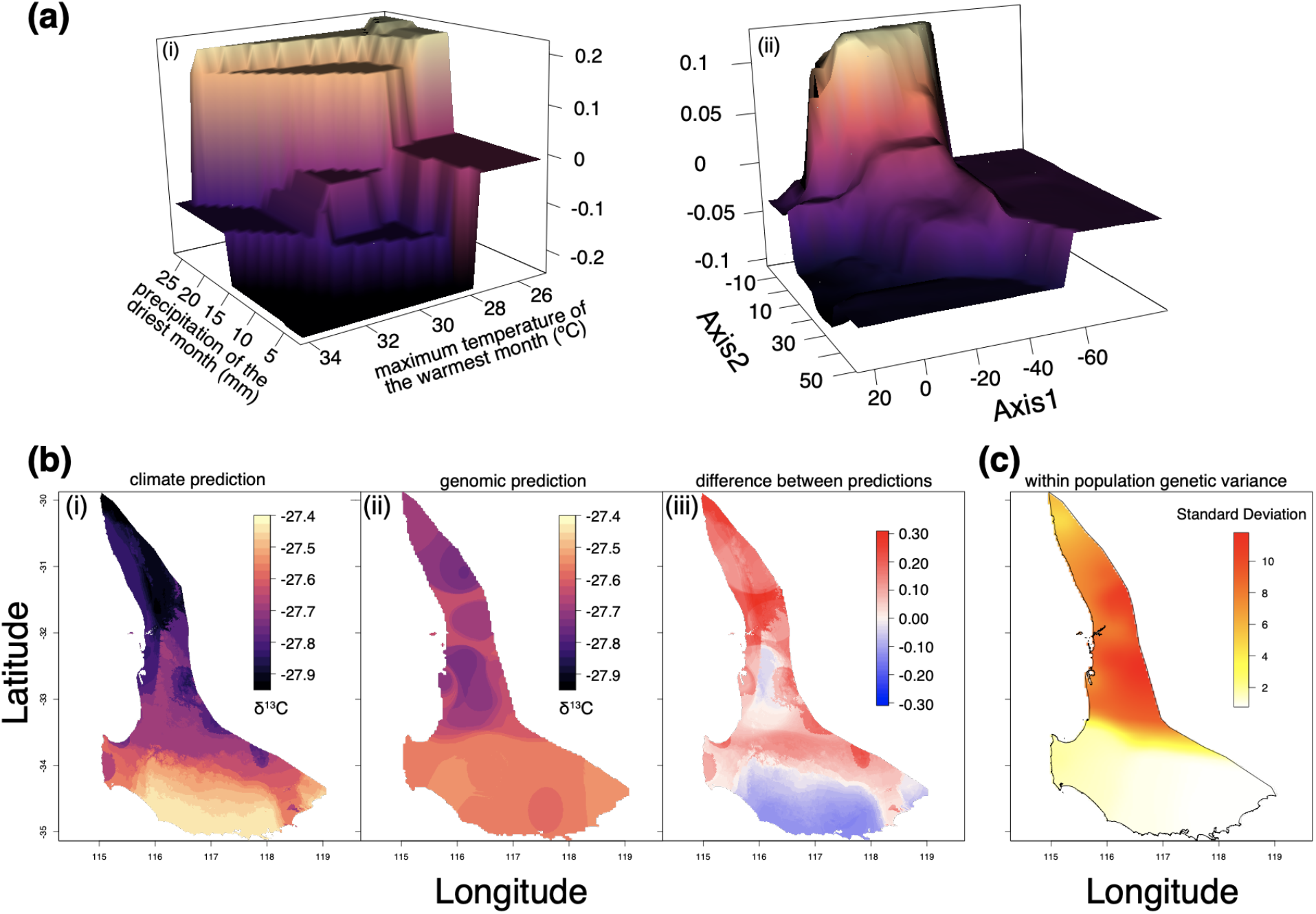
Random forest model of variation in δ^13^C and climate for *Corymbia calophylla.* (a) Predictive landscape 3D surface plot for climate variables predicting δ^13^C (a.i.) and candidate SNPs predicting δ^13^C (a.ii.). (b) Predictions across the distribution for climate (b.i.), candidate SNPs (b.ii.), and the difference between them (b.iii.). (c) The genetic variation measured as standard deviation within populations among candidate SNPs associated with δ^13^C.

These maps are only indicative of the variance associated with trait means and are only partially indicative of the patterns of adaptive capacity within natural populations; therefore, we explored the distribution of population-level adaptive genomic variance by plotting the additive standard deviation from both DAPC axes (Figure 4c). We found that the populations in the northern portion of the distribution have more adaptive variance than their far north or southern counterparts, but the lack of variation is particularly evident in the southern region. This suggests that as the climate continues to change, the southern region will be at high risk of decline because it lacks the adaptability associated with drought related standing genetic variation needed to persist in future environments. We compared this variance map to a similar map from associations with precipitation of the driest month (Figure S13 & S14) and found that the variation associated with climate is high in one population and failed to identify critical variation in the greater northern region associated with drought traits, underpredicting the variation. In total, we found that within-population adaptive variance changes through the landscape and is likely critical for the persistence of the species, a finding that is consistent with those in Anderegg et al (29) that trait variation is likely more important than trait means for adaptation to new climate challenges. These findings challenge current approaches that use climate to predict future genetic or trait distributions, because genotype-environment associations or trait-climate predictions, by design, are limited in identifying within-population variation as adaptive at the ends of the climate spectrum. This is likely due to the fact that traits are polygenic, additive, and adapt to climates in uneven or skewed ways.

*Corymbia calophylla* possesses heritable traits important for adaptation to drought related events and the mean trait distribution shows clear patterns of association with climate (17, 30). Our spatial predictions from genomic associations are more constrained than climate predictions. Yet, genomic predictions had greater importance values than climate predictions (δ^13^C = 32.9_GENO_ & 25.0_CLIM_; NDVI = 0.19_GENO_ & 0.13_CLIM_; SLA = 11.2_GENO_ & 10.1_CLIM_), but lower variation explained (δ^13^C = 53%_GENO_ & 69%_CLIM_; NDVI = 21%_GENO_ & 49%_CLIM_; SLA = 35%_GENO_ & 81%_CLIM_). This difference might be because climate includes non-heritable trait variation, might compile additive variation due to small-effect alleles, or may capture the overall effects of epistatic interactions. In this way, the climate-only models may overpredict spatial distribution of trait values because it includes non-heritable variation, and it is also unable to ascribe where the drought variation occurs. While genomic prediction models may underpredict spatial distribution of trait values. Importantly, we show that northern populations have greater standing genetic variation associated with drought tolerance traits, while southern populations will likely be deficient in adaptive variation. The adaptively deficient populations will require receipt of adaptive variation from outside sources, either through natural gene flow and pursuant selection or through human-mediated gene transfer such as assisted gene migration or climate adjusted provenancing (31).

## *Cis*-regulatory variants drive trait adaptation

The adaptive variation associated with traits, particularly for δ^13^C, is largely driven by variants in *cis*-regulatory regions (noncoding DNA that regulates neighboring genes), which are less constrained by pleiotropy than coding regions from an evolutionary perspective (32); and this mechanism appears to be important in *C. calophylla*. Indeed, recent studies suggest that *cis*-regulatory regions are critical for different types of adaptation (33–35). Yet there is poor understanding how this variation influences population-level local adaptation, as noted by recent studies on evolution (36). Here, we characterise variants associated with functional traits that are important for this species’ adaptation to climate in *C. calophylla* that are overrepresented by *cis*-regulatory regions. This finding suggests that adaptation within *cis*-regulatory regions are more abundant than variants found within protein-coding genes and are more likely to shape the genomic architecture of local adaptation for these leaf-level traits. Similarly, parallel evolution of loss of flight in birds has been associated more strongly with non-coding regulatory DNA than in protein-coding genes (37). We therefore contend that, similar to the evolution of interspecific phenotypes being largely driven by change in regulatory regions, adaptation of intraspecific phenotypes to different climate exposure is also governed by changes within regulatory regions.

## Conclusion

We assessed the genomic contribution to trait variation and found that relatively few variants explained a large proportion of the trait variation. We also observed that the genomic variation associated with traits was aggregated in a few genomic regions. It appears that adaptive genomic regions are important for phenotypic evolution, and it has been postulated that these regions may facilitate rapid phenotypic divergence, such as that observed in drought tolerance in maize (38) and parallel evolution among freshwater and oceanic populations of three-spine stickleback (39).

Elucidating intraspecific patterns of local adaptation to contrasting climate has been an important objective of evolutionary biologists for decades, leading to outcomes aimed at improved management and conservation of species in future climates. This is especially true for ecologically important tree species that drive ecosystem interactions, such that loss of the species leads to the decline in other species due to direct and indirect dependencies. Our results integrate genomic, climatic, and phenotypic data, within a genetically diverse non-model system, to illustrate the complexity of trait determination. Notably, we found that standing genetic variation associated with trait variation occurred within small, gene-rich windows, and that this variation was variable across the species range such that within-population variation may be more important than species-wide variation for adaptation to the effects of climate change. We challenge current dogma that populations most adapted to the driest temperatures are the best sources for adaptive variation. We show that if we choose these populations for assisted gene migration based on adaptation to drought, then the gene pool would be depauperate of genetic variation associated with important traits, rather by using populations that are both adapted to their current conditions and adaptable to more extreme conditions, new populations would have enhanced adaptability to novel local conditions through new patterns of recombination and the emergence of beneficial trait values. However, we must bear in mind that managing complex traits is not so simple because epistatic and pleiotropic interactions among candidate alleles are frequent and may complicate ongoing management strategies, i.e., by promoting one trait or gene, another trait or gene may be unexpectedly promoted or suppressed.

We also show that modeling the distribution of drought response traits changed depending on the use of climate or SNPs, and that climate-only models may have over-predicted population-level adaptive potential. Ultimately, genomic predictions of important physiological traits for an ecologically important species provides improved understanding of the complexities of climate adaptation and provides a way in which we might harness that information to protect forests at their physiological limit from changing climate conditions.

## Methods and Materials

### Study species

*Corymbia calophylla* is a foundation forest canopy species located in Western Australia (WA). It is considered a foundation species because it is critical for forest structure and ecological processes (1). This species is an ideal candidate in which to study adaptation of functional traits because its distribution traverses strong environmental gradients over short distances, it has recently experienced mortality events attributed to climate change (2, 3), and evidence of adaptation to climate has been identified in physiological experiments and genome-environment investigations (4–7).

### Experimental site

This research was conducted in a plantation near Margaret River, WA Australia (Figure 1 main text), located in the *C. calophylla’s* cool-wet region. Seed collection and trial design are described in detail in Ahrens et al. (2019). Briefly, 18 populations represented by 165 families were established at the experimental site for a total of 3,960 individuals in six replicated blocks with two rows of buffer trees to minimise edge-effects. Families are defined here as individuals that have a known, common mother but unknown fathers (i.e., half-sibs) via mixed pollination within an intact forest. We focused on 12 populations representing contrasting climate combinations covering the full geographic distribution of C. calophylla (Figure 1 main text). We sampled phenotypes and genotypes from a total of 432 trees, including 4 half-sibs from 10 families within 12 populations when available for a total of 120 families.

### Trait measurements

Traits were measured in March 2017 on *C. calophylla* trees that were 29 months old and 2-3 m tall. For each individual tree, we removed a north facing, mid-canopy side branch at its intersection with the main stem. The side branch was removed in the morning (between 8 a.m. and 12 noon), stored in a cool box, and measured in the afternoon (between 12 noon and 6 p.m.). For each side branch, we collected data for the three traits (among others not listed here): integrated water-use efficiency (δ^13^C), specific leaf area (SLA), and normalized difference vegetation index (NDVI). All traits have shown close association to climate in past studies. High water-use efficiency (WUE) is the link between photosynthesis and evaporation (8) that translates to climatic tolerance under water limitation. Water-use efficiency is correlated with isotope discrimination (δ^13^C, an isotopic signature measuring the ratio of 13C and 12C (9)) and relates to leaf gas exchange properties (10, 11). To estimate δ^13^C, the leaves were kept in an airtight box with silica gel until they could be dried in an oven at 70°C for 48 hr. δ^13^C was measured from leaves dried using a benchtop freeze dryer (Alpha 1-4 LDplus Laboratory Freeze Dryer, Martin Christ). The leaves were grounded into a fine powder using a cyclotec mill (Foss Analytics) and sent for isotope analysis (ANU Isotope Laboratory) using a coupled EA-MS system (EA 1110 Carlo Erba; Micromass Isochrom).

Leaf-level normalized difference vegetation index (NDVI), which is generally used to measure chlorophyll content by quantifying leaf greenness, and is closely related to fraction of absorbed photosynthetically active radiation (FPAR) (12, 13). While not technically a functional trait (NDVI), traits based on spectral properties of leaves can be indicative of photosynthetic activity and plant stress, and from hereon, we include this complex trait as a functional trait for ease of discussion. A field spectroradiometer (ASD standard-resolution FieldSpec4, Malvern Panalytical) was used to measure leaf reflectance in the visible and reflected infrared spectral regions with 2,151 narrow bands (10 nm full width at half maximum) and 1 nm spacing between band centers. Measurements were made for three leaves using a leaf-clip attachment with its own light source and calibrated to % reflectance using data collected from a Spectralon white reference panel. Means for all bands among the three leaves were calculated for each individual tree. Specific wavelengths were used to estimate the modified red-edge NDVI. The modified red-edge NDVI was calculated using the following equation (14):

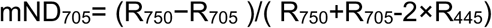

and was developed as an improvement to the standard NDVI to provide a more robust estimate of chlorophyll content (15) across a wide range of species and leaf structures (14). Henceforth, this index will be referred to as “NDVI” in the text.

Specific leaf area (SLA) varies across global climate gradients (16), and high SLA values increase tree susceptibility to drought-induced mortality (17). Specific leaf area (SLA) was measured on three fully matured leaves that were representative of the branch. After removing half of the petiole with a razor, the leaves were scanned into a computer using a Canon flatbed scanner (model # LiDE220) at 50 dpi. The leaves were then dried in an oven at 70°C for 48 hr and leaf mass was estimated on a digital scale with 1000th of a gram accuracy. SLA was calculated by dividing total leaf area by total leaf mass for all three leaves and averaged across the three leaves for a single SLA value for each individual tree.

### BLUP estimation

Best linear unbiased predictions (BLUP) were estimated for each trait to account for variation attributed to the design matrix and to increase trait accuracy because it anticipates regression of progeny to the mean observed (18). Analysis was performed using ASreml Version 4.1 (19, 20). Univariate variances were estimated within the framework of the linear mixed model:

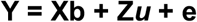

Where **Y** is the column vector of individual phenotypic values of the response variable, **X** is the design matrix associating observations with fixed effects, ***b*** is the vector of fixed effects, **Z** is the design matrix associating observations with random effects, ***u*** is a vector of random effects and **e** is the vector of residual errors assumed to be identically and independently normally distributed with *E*(*e*) = 0.

Two sets of analysis were conducted, the first at the family level for the purpose of checking the data for homoscedasticity and determining if there was a need for transformation, and the second at the individual tree level for the purpose of estimating BLUPs.

#### Univariate family model

Elements in ***b*** included the intercept and provenance effects while elements in ***u*** included replicate, row within replicate, column within replicate, plot and family. Residual plots were examined for homoscedasticity and appropriate transformations identified as outlined by Gilmour et al. (2002). The trait NDVI was log transformed, whereas the δ^13^C and SLA did not require transformation.

#### Univariate individual tree model

Elements in ***b*** and ***u*** were the same as for the univariate family model, with the exception that the family term was substituted with an individual tree, random additive effect. In this model, additive genetic covariance between relatives is modelled via the numerator relationship matrix (Henderson 1976). A one-tailed log likelihood ratio test with 0.5 degrees of freedom (20, 21) was used to test the significance of additive variance estimates for each trait.

### Reference genome

#### DNA extraction and sequencing

We isolated high molecular weight DNA suitable for long-read re-sequencing by following a nuclei and magnetic bead-based extraction protocol (Jones and Borevitz, 2019). Briefly, 30 g of fresh leaf material from an individual from the Australian Botanic Gardens in Canberra Australia was processed with 150 ml nuclei isolation buffer using a high-powered blender. The homogenate was filtered using a funnel of Miracloth. Next, 100% Triton X-100 was added to extract the nuclei from chloroplasts. The nuclei pellet was washed twice with a chilled nuclei buffer. Nuclei pellet lysis was performed with a lysis buffer containing 3% Sodium dodecyl sulphate (SDS) followed by incubating at 50°C. The DNA was cleaned of proteins by adding potassium acetate and pelleting. The supernatant was bound to Sera-MagTM SpeedBead magnetic carboxylate-modified particles (GE Healthcare). The beads were washed with 70% ethanol until clean. Size selection for fragments ≥30 kb was performed using a PippinHT (Sage Science, Beverly MA). MinION Mk1B was used to sequence the long-reads ( Oxford Nanopore Technologies, ONT).

#### Nuclear genome assembly

Raw read libraries were filtered and trimmed in preparation of assembly with NanoPack ((22); NanoLyse version 1.1.0; NanoFilt version 2.6.0). First, ONT DNA control strand was removed. Next, 200 bp was trimmed from both 5’ and 3’ ends, removing sequencing adapters and low quality read ends. Finally, filtering removed all reads less than an average quality of 7 and less than 1 Kbp in length. Quality controlled read libraries were de novo assembled using the long read assembler Canu ((23); version 1.9; parameters: corOutCoverage = 200 “batOptions = −dg 3 −db 3 −dr 1 −ca 500 −cp 50”, correctedErrorRate = 0.154, corMaxEvidenceErate = 0.15, -fast). Following assembly, contaminant contigs were identified with blastn ((24); version 2.9.0+) using the NCBI nucleotide database (NCBI Resource Coordinators, 2013; versions BLASTDBv5). Identified contaminant contigs were removed with Blobtools (25); version 1.1.1). Haplotigs, assembly artifacts and plastid contigs were removed from assemblies with purge haplotigs (26); version 1.1.0). Next, all assemblies were polished with the long read polisher Racon (27); version 1.4.11) combined with minimap2 and the short read polisher Pilon (28); version 1.23) combined with BWA-MEM ((29)). Contigs of less than 1 Kbp were removed and manual curation of all remaining contigs was performed with MUMmer (30); version 4.0.0beta2) to identify plastid DNA. Finally, our assemblies were scaffolded by RaGOO (31)) using synteny information provided by the previously published *Eucalyptus grandis* genome (32). Genome completeness was assessed with BUSCO ((33); version 3.0.2) and Lai ((34): version beta3.2). See figure S2 for a summary of our assembly statistics.

#### Chloroplast Assembly

A chloroplast genome was assembled using the curated ONT libraries. Chloroplast reads were identified and extracted by aligning all reads against a composite chloroplast genome made up of all published *Eucalyptus* genomes. Identified chloroplast reads were assembled with Unicycler (35); version 0.4.8) and polished with pilon.

#### Repeat and gene annotation

Prior to gene annotation repetitive regions in *C. calophylla’s* genome were identified and soft masked with RepeatMasker (36); version 4.1.1) using *de novo* repeat libraries created with EDTA ((37); version 1.9.6). Protein-coding genes and transcripts were predicted by BRAKER2 (38); version 2.1.5; parameters: epmode), using proteins sequences for Myrtaceae (Taxonomy ID: 3931) and Arabidopsis thaliana (Taxonomy ID: 3702) obtained from the National Center for Biotechnology Information (39) as homology evidence.

### Library preparation & variant calling

DNA extraction for whole genome sequencing was performed by the Australian Genomic Research Facility (AGRF, Adelaide, SA Australia) using a modified ctab method. We generated short-read whole-genome shotgun DNA sequencing libraries using a low cost transposase-based protocol (40). Briefly, we quantified DNA concentrations using a fluorometric Quant-iT™ high sensitivity dsDNA assay kit (Molecular Probes™ Q33120). To normalise concentrations among samples, we diluted DNA to 2 ng/μl, quantified again and then diluted to 0.8 ng/μl. To form sequencing libraries, we combined 3 μl of each sample (approx 2.24 ng) with a small quantity of a Nextera™ tagment DNA enzyme (Illumina catalogue #15027865). To decrease costs, we performed this tagmentation reaction at 1/5th volume and 1/5th concentration of the manufacturer’s protocol, i.e. 1/25th reactions. We amplified the libraries and added custom index sequences during 13 cycles of PCR (primer sequences provided in (40)). We purified and size-selected libraries using two SPRI-bead based cleanups and electrophoresis-based final size selection for insert sizes between 200 and 500 bp. We sequenced these libraries on a single S4 flow-cell on an Illumina NovoSeq 6000 instrument at Genomics West Australia/Telethon Kids Institute, Perth, West Australia.

Sequencing yielded between 3 Gbp and 10 Gbp per sample (~10-30X coverage), pooled across all sequencing runs (see Figure 2 in main text). We discovered genetic variation among samples following the approach of Murray et al. (41). Briefly, we filtered, trimmed, and merged pairs of raw sequencing data using AdapterRemoval (Schubert, Lindgreen, & Orlando, 2016), then aligned reads to reference genomes using BWA-MEM version 0.7.15 (29, 42). We detected short genomic variants using bcftools mpileup (43), normalised variants with bcftools norm (43), and performed initial variant filtering with bcftools filter (43). Reads were aligned against our custom *Corymbia calophylla* reference genome. During initial variant filtering, we discarded variants with quality <10, fewer than five reads in total across all alleles in all samples and fewer than three reads supporting the alternate allele across all samples. Resulting in 91 million SNPs, a SNP every ~4bp.

### Filtering

After variants were called using the above pipeline, additional filtering was performed in PLINK 2.0 (44) with the following thresholds. Minimum read-depth was set to six. We extracted biallelic variants only, to ensure all variants were biallelic and minimise complex signals. The minimum basepair distance between variants was set to 10. Minor allele frequency (MAF) was set to 0.01, to have sufficient power for GWAS detection. Missing data threshold was set to 0.5 but the average missing data in the data set was 0.2. Resulting in a dataset with 6.5 million SNPs across all 11 chromosomes.

### Linkage disequilibrium

Linkage disequilibrium (LD) was measured using median base pair distance to half-maximal *r*^2^ values using boringLD v0.3.0 (https://github.com/kdmurray91/boringld). We set the window size to 30 kbp with a 15 kbp overlap. ts. We fitted analytical models of the decay of *r*^2^ as a function of inter-SNP base pair distance using formulae derived by Hill and Weir (1988) and then calculated base pair distance to half-maximal *r*^2^ for each window. We summarized per-window esti-mates of half-maximal *r*^2^ across all genome windows for a global *r*^2^ estimate. To test if LD was a function of the number of SNPs within each window, we used a linear model within each chromosome (Figure S15). The linear fit was significant for all chromosomes but the *r*^2^ values were low, this pattern was driven by the windows with very few SNPs.

### Associations

Genome wide association studies (GWAS) were performed in Plink2 for each of the three functional traits. We used the individual BLUP estimates as the functional trait inputs, as this accounted for experimental site effects. We used the first 10 axes from an MDS as a covariate for population structure. We also tested the associations of the GWAS using the traditional approach with a kinship matrix as a random variable in GEMMA and the outputs resulted in similar significant variants to the Plink2 analysis, and we therefore focus on Plink2 outputs. We used the general linear model (glm) function to calculate *p*-values. We imported the Plink2 results to R and adjusted the p-values for multiple comparisons using the Benjamini-Hochberg method. Then used the *CMplot* command from the *CMplot* package to visualise the *p*-value distribution in a manhattan plot format. To ensure that these signatures of selection identified in Plink2 were not random, we permuted the phenotype data in two ways and ran the Plink2 GLM analysis for both permutations. The first was completely random using the sample function in R without replacement (Figure S16b-S18b), and the second was keeping the family structure of phenotypes and resampling among families (Figure S16c-S18c).

The qq plots suggest that the *p*-values were inflated when controlling for population structure (PLINK) and under-dispersed (uniform distribution) when controlling for family structure using a kinship matrix (GEMMA; Figure S19). Both GWAS methods identified very similar groupings of SNPs (51-65% total similarity and top 50 SNPs were 88% similar for δ^13^C). The differences between the distribution of BH adjusted *p*-values were significantly different between the real and permuted datasets (t.test: T_1,1.3m_ = −2648.8, versus Random *p* < 0.001; T_1,1.3m_ = −244.4, versus Family p < 0.001), suggesting that the adaptive variants were not due to chance. We wanted to ensure that the associations between SNP and trait were associated with local genomic structure, as described in Li and Ralph (45). Therefore we used the package lostruct in R to investigate the structure within 1000 SNP windows (this is equivalent to approximately 10 kbp) within each chromosome, creating between 400 and 600 windows per chromosome. Then we compared the first two axes within chromosomes to the location of adaptive genomic regions for three major areas of association on chromosome 3, 8, and 10. We found that anomalous local population structure among 1000 SNP windows was not localised near regions that were significantly associated with phenotypes (Figure S20).

### SNP Heritability

In order to calculate SNP based heritability, we used the GEMMA model to describe the proportion of variance in phenotypes explained (PVE). GEMMA fits a univariate linear mixed model for marker association tests with a single phenotype, and for estimating the PVE by all variants (46).

### Genomic structure of SNPs associated with phenotype

For the 279, 69, and 92 SNPs that were deemed significantly associated with the three traits, we generated discriminant analysis of principal components (DAPC) plots and recorded the eigenvalues as a means of describing genomic differentiation at the cluster level. Indeed, DAPC is useful here because it optimises variance among groups while minimising variance within groups. We identified the number of clusters using the *find.clusters* argument in the adegenet package in R. Our goal here was to explain as much variance as possible by using as few PCAs as possible. Therefore, we kept 5, 7, and 3 PCA axes for δ^13^C, SLA, and NDVI respectively, while identifying 7, 7, and 4 genomic clusters among the 12 populations. The variation explained for each axis was calculated by dividing the eigenvalue form the first or second axis by the sum of all eigenvalues. The DAPC coordinates for each axis were recorded for input into the random forest model described below in the ‘predicting trait distribution’ section.

### Epistasis & pleiotropy

We attempted to uncover some of the complex epistatic and pleiotropic relationships between variants and traits by use of the combined analysis of pleiotropy and epistasis (CAPE) package in R (47), which implements an analytical method described in Carter et al. (48) to explicitly test for these complex interactions. This method was designed for datasets that include populations with mixed genetic variation, and is therefore appropriate for our study design. CAPE calculates both the main effects, which are the effect of a SNP from the set of all pairwise regressions that included that SNP, and the directional influences of that SNP that interact epistatically. We used the following parameters for the CAPE analysis (parameter file available online): traits_scaled = true, pval_correction = fdr, alpha = 0.5, peak_density = 0.8, tolerance = 10, num_alleles_in_pairscan = 300, maxpair_cor = 0.5, pairscan_null_size = 1000. We used a high peak_density because of the quick LD decay, as suggested in the CAPE documentation. We also used a num_alleles_in_pairscan of 300 to limit the number of SNP pair analyses, this results in a different outcome for each run because we do not test all 104,329 pair possibilities. To be clear, individual SNP pair outcomes will not change, it is whether or not the individual SNP pair is randomly included in the output. Even so, the result shown here is a representative subset of these interactive effects. Both the inputs and outputs for our specific CAPE analysis are provided online, so the user can recreate our figures but also explore other individual runs and create new figures.

### Predicting trait distribution

To predict how climate and significantly associated SNPs independently predict trait BLUPS, we used random forests (49) implemented in the extendedForest software package v1.6.1 in R, built upon the original randomForest package (50). We chose to use the extendedForest package because it uses a modified version of calculating variable importance. Permutation importance is calculated by passing the out-of-box (OOB) samples reconfigured with the extendedForest permutation grid through each respective tree in the standard way, and bootstrapped with two-thirds of the data to construct a tree. A regression tree is then fitted to the bootstrapped data, where each node split is based on a subset of M predictor variables. This procedure is repeated B times, where B is the number of trees grown. If a large number of trees are grown (e.g. > 1000), then the generalisation error (e.g. the true error of the population) is minimised such that overfitting of the training data is not possible (51, 52). The accuracies and error rates of each tree are computed using the OOB sample. As the OOB sample is withheld for the tree growing procedure, the OOB estimates are essentially equivalent to a N-fold cross-validation (51). An advantage of the OOB sample is that the use of OOB error estimates removes the need for a test data set, and the estimates of error are said to be unbiased (49). We created two models for each trait, the first with the first two DAPC axes and a second with the two most important climate variables. Climate variables were chosen based on previous knowledge of the species and traits (aridity index, potential evapotranspiration, maximum temperature of the warmest month, minimum temperature of the coldest month, precipitation of the driest month). We used the following parameters for all random forest models: sample size = 320, number of trees = 5000, mtry = 2 (optimal number of predictor variables), nperm = 100, keep.inbag = true. For climate-only models, the random forest model was repeated for the top two most explanatory climate variables for each, and this output was used for further exploration. Additionally, random forest uses the OOB samples to construct a variable importance measure that describes the relative strength of each predictor variable.

While the multidimensionality of this function is difficult to visualise, two predictor variables can be viewed using simple line plots (e.g. univariate partial dependency plot) or perspective mesh plots (e.g. bivariate partial dependency plots). We used the forestFloor package to create univariate line plots for each variable and 3d mesh plots using the plot and show3d functions in the forestFloor package in R to describe the nonlinear relationships between dependent and independent variables (53). We then wanted to visualise how the predictions were distributed in geographic space, therefore, we used the predict function between a raster stack of the two explanatory variables and the results from the random forest model to predict the distribution. Climate rasters were downloaded from worldclim and clipped in R using the raster package (54), while the genomic rasters were created using QGIS based on the 12 populations and interpolated using the kriging method. The resulting predictive map was plotted using the plot function in R using the same trait prediction scale. To visualise the difference between the two predictions we subtracted the climate prediction from the genomic prediction (TR_DIF_ = TR^p^_GEN_ - TR^P^_CLI_) and plotted the map in the same manner as above. Lastly, to visualise which areas have the greatest genomic variability, we calculated the standard deviation among each DAPC axis, mapped them independently, and added the genomic standard deviation maps together for one estimate of genomic variability, and smoothed them using the gstat argument in the package of the same name with the following parameters: nmax = 4 and set = list(idp = .5) (55). The interpolation was done using the interpolate function in the raster package.

### Functional annotations and GO enrichment

The program snpEFF (56) was used to identify the location of significantly associated SNPs using the *Corymbia calophylla* genome (NCBI txid34324; assembly ASM1418284v1). Variants found within genes were recorded as synonymous or nonsynonymous, in addition variants in regulatory regions found within 5,000 base pairs of genes were recorded as being upstream or downstream, along with the number of base pairs between the gene and SNP. We also specified which variants are in promoter regions, defined here as being within 500 bp upstream of the gene. Then, orthofinder was used to identify orthologs between *C. calophylla* and *E. grandis* genomes (57), and assign putative functions to predicted genes identified as significant. We used the *E. grandis* gene IDs to identify within trait higher order functions through gene ontology enrichment using the phytozome website and the *E. grandis* organism. Gene networks of GO enrichment terms for each process and trait were created using revigo (58) and the networks were visualised using Cytoscape v3.8.2 (59).

The 11 genes that were found to be significantly associated with all three traits are expressed during growth and development processes, including in plant structure such as guard cell and leaf structure (Eucgr.A00085, Eucgr.B03379, Eucgr.C00318, Eucgr.K02549, Eucgr.H02572, Eucgr.H02574, Eucgr.H02575 and Eucgr.C00581).

