## supplementary information for "Genomic constraints to drought adaptation"


| Table/Figure # | text | Title | page # |
|---|---|---|---|
| Table S1 | main | Assembly statistics for *C. calophylla*. | 2 |
| Table S2 | main | Linkage disequilibrium summary table. | 3 |
| Table S3 | main | Annotation summary | 4 |
| Figure S1 | main | Linear regressions between raw trait value the BLUP | 5 |
| Figure S2 | main | Summary histograms for the reference genome | 6 |
| Figure S3 | main | Histograms for chromosomal LD | 7 |
| Figure S4 | main | Diagnostic quantile-quantile plots | 8 |
| Figure S5 | main | Manhattan plots for all nine traits | 9 |
| Figure S6 | main | MDS of the first two axes for population structure covariance | 10 |
| Figure S7 | main | LD ($r^2$) between SNPs within trait and among traits | 11 |
| Figure S8 | main | Full variant network output from the cape analysis | 12 |
| Figure S9 | main | Gene networks from GO enrichment analysis | 13 |
| Figure S10 | main | DAPC for each trait | 14 |
| Figure S11 | main | Random forest analysis for NDVI | 15 |
| Figure S12 | main | Random forest analysis for SLA | 16 |
| Figure S13 | main | Manhattan plot for BIO14 | 17 |
| Figure S14 | main | Location of genetic variation for BIO14 | 18 |
| Figure S15 | main | Linear regressions between SNPs and LD | 19 |
| Figure S16 | methods | Trait permutation to test for $\delta^{13}C$ | 20 |
| Figure S17 | methods | Trait permutation to test for NDVI | 21 |
| Figure S18 | methods | Trait permutation to test for SLA | 22 |
| Figure S19 | methods | Image plot of the kinship matrix | 23 |
| Figure S20 | methods | Local genome population structure | 24 |

Table S1. Assembly statistics for *C. calophylla*.

| | Un-scaffolded | | Scaffolded | |
|---|---|---|---|---|
| **Genome assembly statistic** | **Length / % of genome** | **Number** | **Length / % of genome** | **Number** |
| **Total (Mbp)** | 394.86 | 422 | 394.90 | 36 |
| **Longest contig/scaffold (Kbp)** | 7,649.18 | --- | 48,491.33 | --- |
| **N50 (Kbp)** | 1,980.23 | --- | 39,866.58 | --- |
| **ESize** | 2,476,596 | --- | 37,436,745 | --- |
| **Genes** | | | | 42,234 |
| **Simple repeats** | | | 1.15% | --- |
| **Transposons** | | | 34.80% | --- |
| **Chloroplast size (bp)** | | | 158,591 | --- |

Table S2 – Linkage disequilibrium summary table. SE = standard error; SD = standard deviation; windows = the number of 30kbp overlapping windows per chromosome; nsnp_mean = the mean number of SNPs per window; nsnp_max = the maximum number of SNPs per window; nsnp_min = the minimum number of SNPs per window.

| CHR | mean | median | max | min | SE | SD | windows | nsnp_mean | nsnp_max | nsnp_min |
|---|---|---|---|---|---|---|---|---|---|---|
| **1** | 468.385 | 211.5 | 75858 | 3 | 44.953 | 1932.448 | 1856 | 483.248 | 1289 | 2 |
| 2 | 315.865 | 121 | 8710 | 3 | 12.218 | 604.158 | 2481 | 442.117 | 1637 | 2 |
| 3 | 460.888 | 118 | 41687 | 3 | 33.397 | 1716.621 | 2671 | 453.957 | 1596 | 2 |
| 4 | 464.103 | 174 | 28841 | 3 | 33.410 | 1425.703 | 1824 | 502.582 | 1464 | 3 |
| 5 | 288.582 | 81 | 12590 | 3 | 12.398 | 697.158 | 3183 | 488.427 | 1684 | 2 |
| 6 | 455.929 | 200 | 16219 | 3 | 17.011 | 895.644 | 2794 | 448.495 | 1449 | 2 |
| 7 | 345.852 | 88 | 43652 | 3 | 25.285 | 1296.446 | 2655 | 465.772 | 1406 | 2 |
| 8 | 423.481 | 148 | 27543 | 3 | 20.238 | 1147.665 | 3241 | 498.036 | 1883 | 3 |
| 9 | 516.830 | 191 | 35482 | 3 | 37.231 | 1545.422 | 1744 | 456.223 | 1231 | 2 |
| 10 | 597.287 | 230 | 28184 | 3 | 32.752 | 1395.707 | 1834 | 448.543 | 1453 | 4 |
| 11 | 740.037 | 214 | 338845 | 3 | 184.084 | 8064.035 | 1937 | 451.173 | 1513 | 2 |

Table S3. Annotation summary at the chromosome level. Ts/Tv = transition vs transversion ratio; syn = synonymous; NS = nonsynonomous; up = upstream, within 5kbp of a gene; down = downstream, within 5kbp of a gene; high = high effect allele; mod = moderate effect allele; low = low effect allele.

| Chr | Length | Variants | rate | Ts/Tv | syn | NS | up | down | high | mod | low |
|---|---|---|---|---|---|---|---|---|---|---|---|
| 1 | 27,855,297 | 477,415 | 58 | 2.69 | 33,140 | 31,195 | 275,030 | 301,861 | 1,133 | 31,195 | 35,973 |
| 2 | 37,224,018 | 581,410 | 64 | 2.68 | 41,212 | 39,269 | 347,667 | 381,326 | 1,541 | 39,269 | 44,792 |
| 3 | 40,088,764 | 638,500 | 62 | 2.83 | 36,473 | 36,223 | 297,589 | 338,288 | 1,479 | 36,223 | 39,573 |
| 4 | 27,387,958 | 486,376 | 56 | 2.69 | 29,489 | 28,267 | 253,848 | 278,156 | 1,132 | 28,267 | 31,964 |
| 5 | 47,765,057 | 820,982 | 58 | 2.83 | 45,485 | 46,371 | 400,197 | 433,848 | 1,949 | 46,371 | 49,160 |
| 6 | 41,948,806 | 665,206 | 63 | 2.50 | 50,266 | 46,334 | 415,792 | 456,759 | 1,654 | 46,334 | 54,516 |
| 7 | 39,866,576 | 650,378 | 61 | 2.82 | 37,187 | 37,840 | 317,198 | 349,265 | 1,631 | 37,840 | 40,367 |
| 8 | 48,491,325 | 853,957 | 56 | 2.73 | 47,288 | 51,348 | 4,249,308 | 4,252,395 | 3,618 | 51,348 | 51,653 |
| 9 | 26,074,809 | 421,748 | 61 | 2.65 | 24,555 | 26,113 | 244,540 | 260,323 | 1,622 | 26,113 | 26,726 |
| 10 | 27,534,854 | 437,473 | 62 | 2.52 | 34,601 | 31,818 | 277,415 | 303,576 | 1,174 | 31,818 | 37,723 |
| 11 | 29,061,480 | 464,194 | 62 | 2.52 | 35,582 | 32,815 | 306,004 | 340,638 | 1,194 | 32,815 | 38,557 |

Figure S1 – Linear regressions between raw trait value (x-axis) the best linear unbiased prediction (BLUP). Traits are $\delta^{13}C$, SLA (specific leaf area), and NDVI (normalised differential vegetation index).

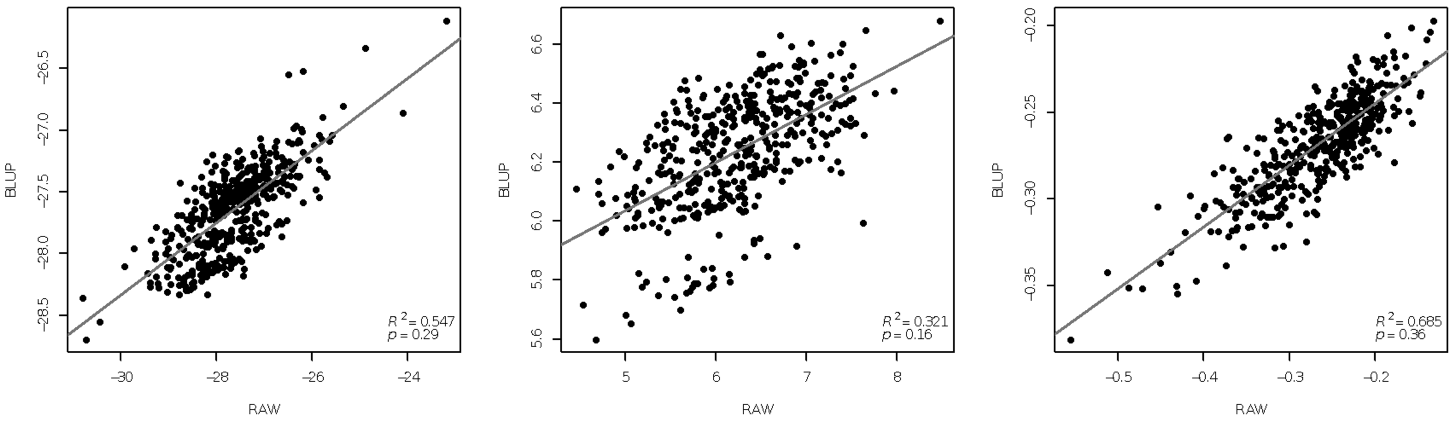

Figure S2. Summary histograms for the reference genome. Top is a histogram of read length. Middle is the filtered and haplotig purged assembly. The bottom histogram is the final scaffolded genome. Showing N50 (orange) and E-size (red) for each histogram.

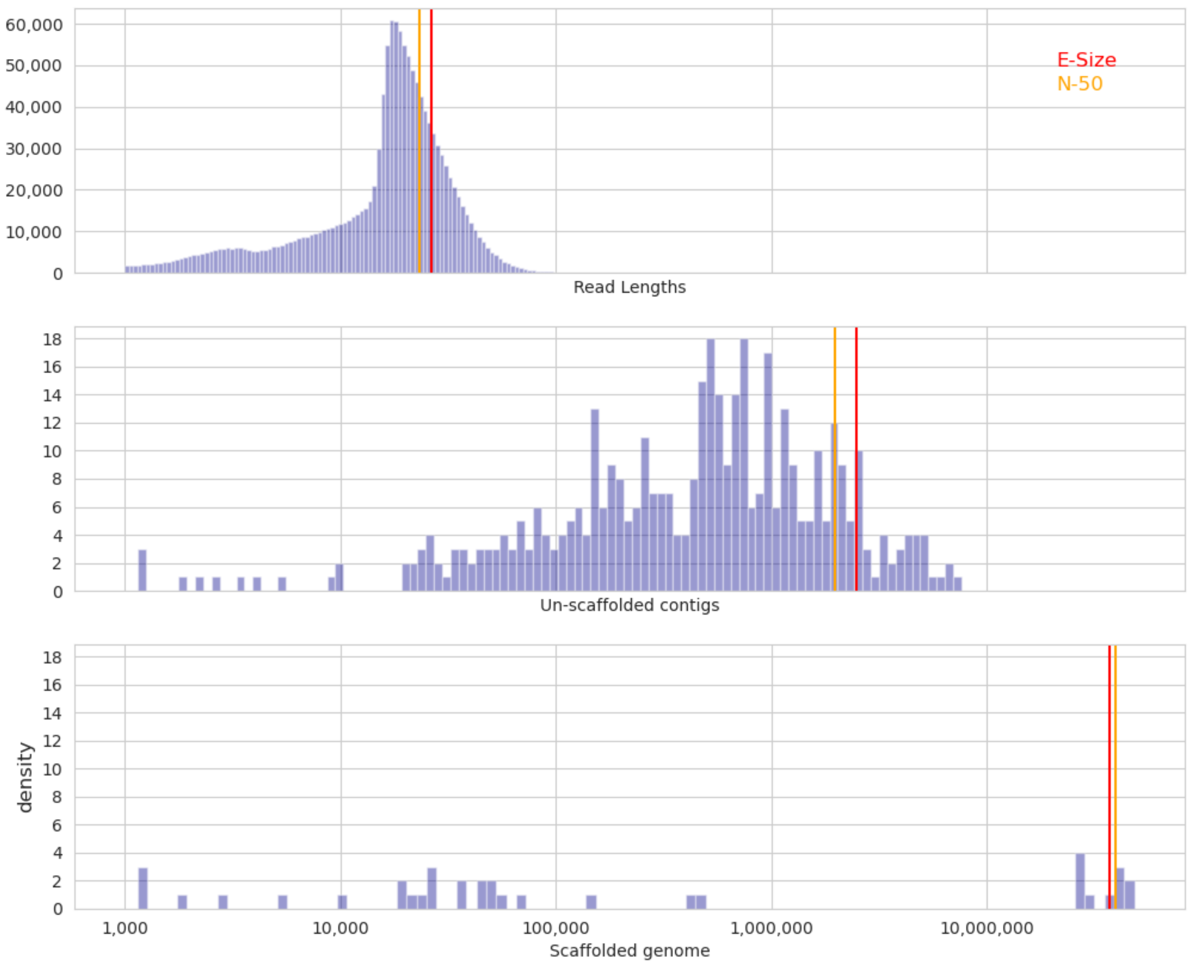

Figure S3 – Histograms for the halfmax calculations that were used to estimate linkage disequilibrium for each chromosome.

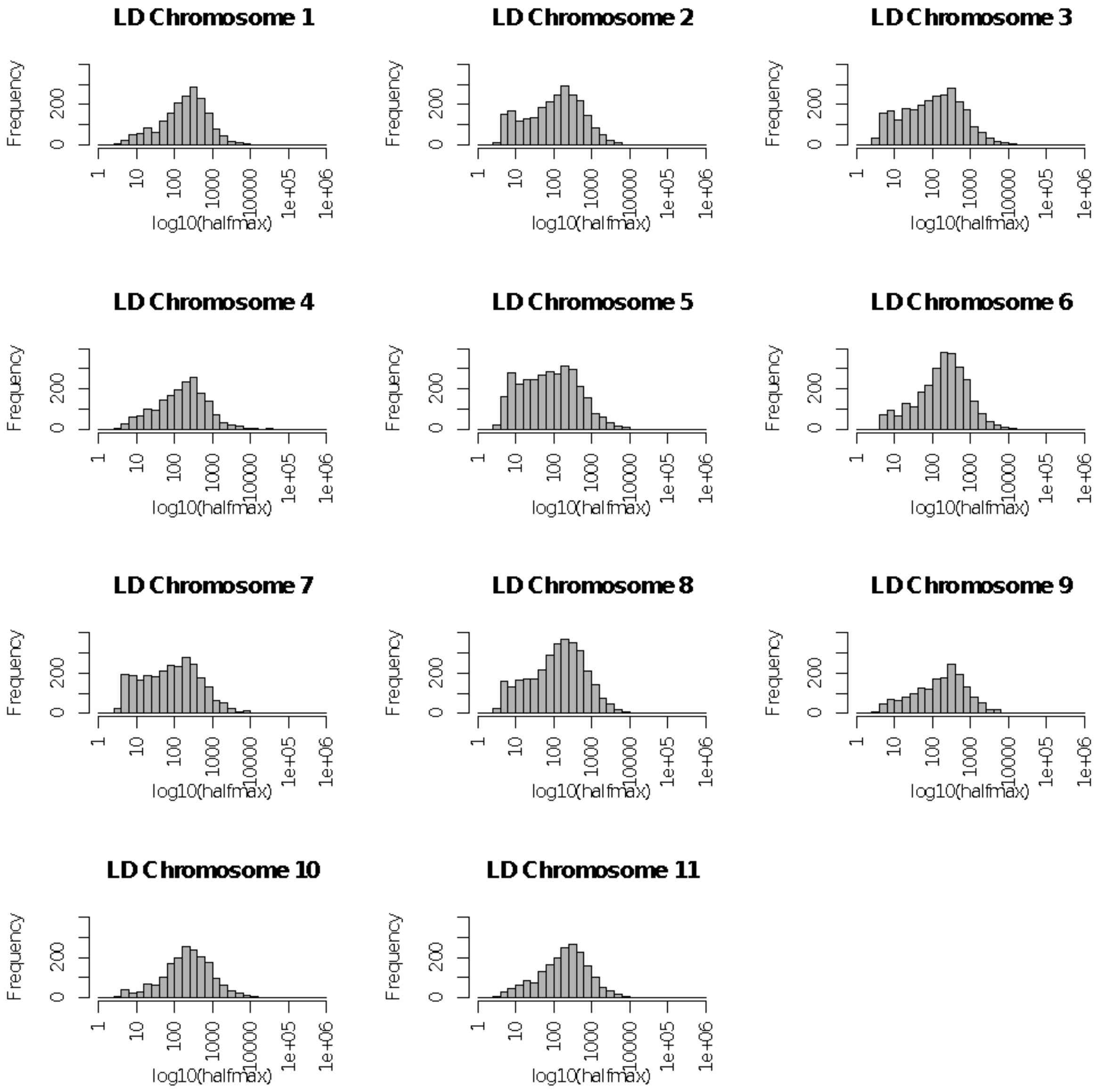

Figure S4 – Diagnostic quantile-quantile ("qq") plot. The observed *P*-value for all SNPs on the *y*-axis versus the expected uniform distribution of *P*-values under the null hypothesis of no association on the *x*-axis. These are the qqplots generated from plink and population structure.

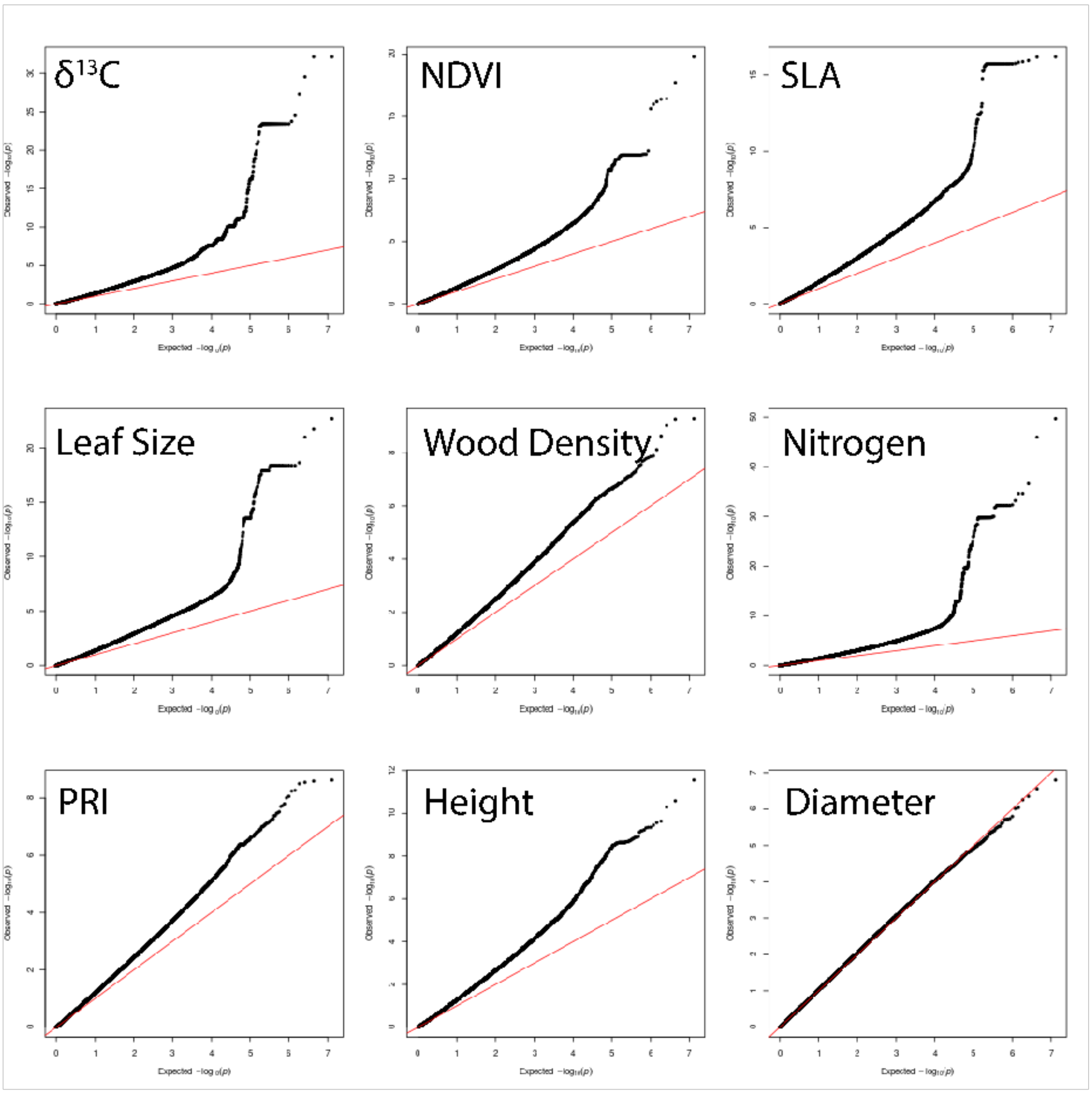

Figure S5. Manhattan plots showing the -log10(adjusted *P*-value) for all nine traits. traits are in this order: $\delta^{13}C$, NDVI, SLA, leaf size, wood density, nitrogen content, photochemical reflectance index, height, and diameter.

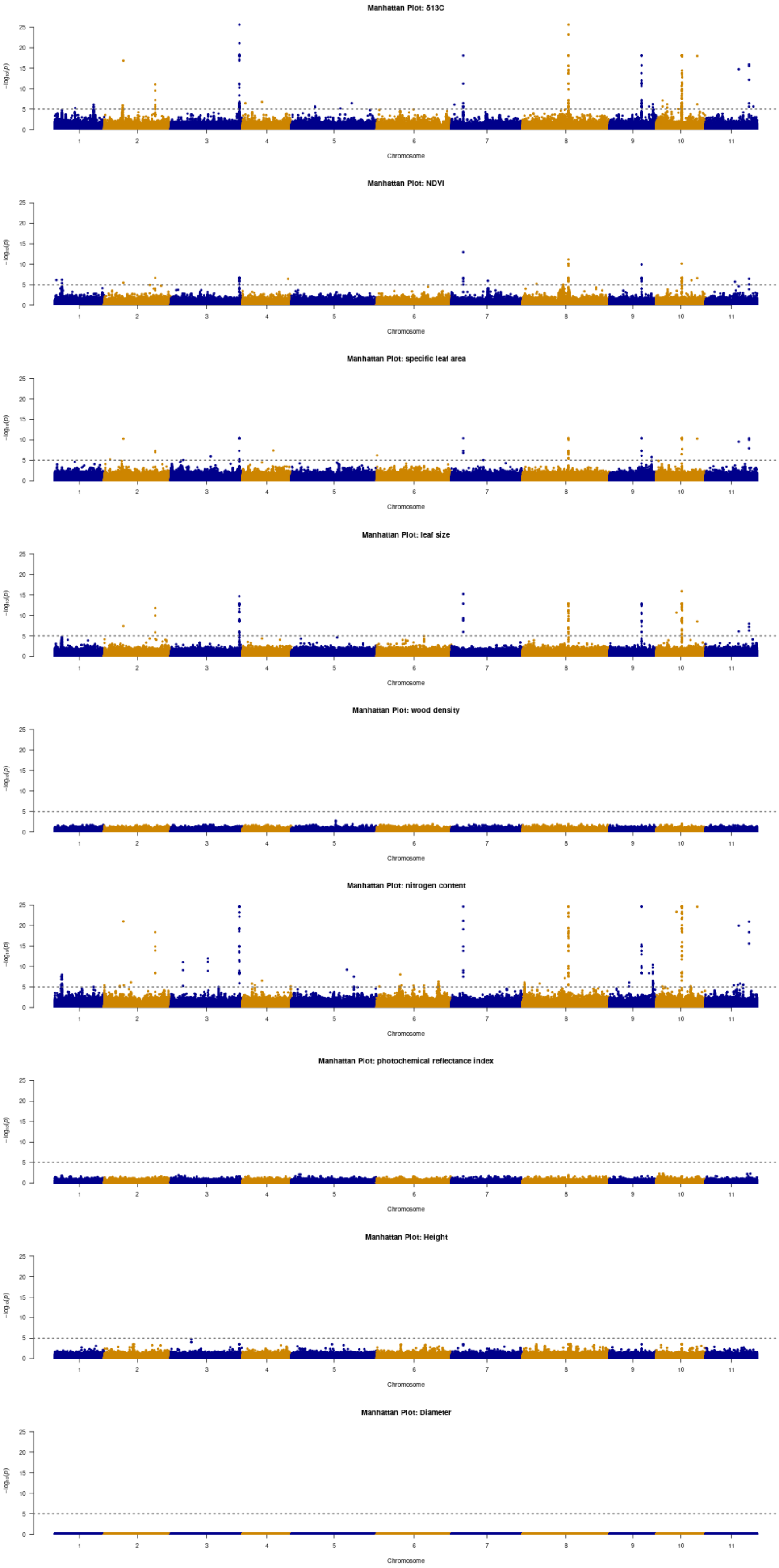

Figure S6 – Multidimensional scaling of the first two axes for covariance estimate in the GWAS analysis. Coloured by populations.

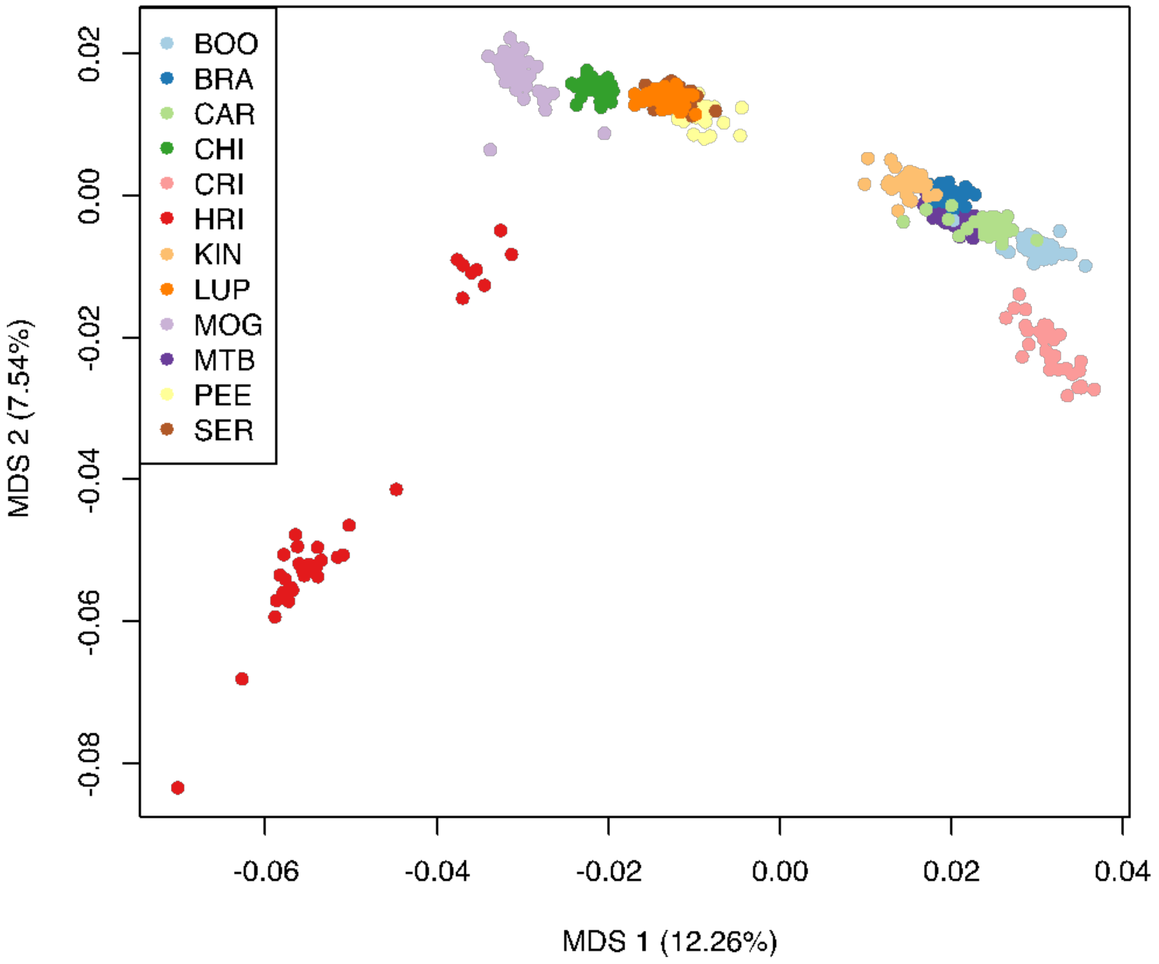

Figure S7. Linkage disequilibrium ($r^2$) between SNPs within trait (a-c), and among traits within chromosomes (d-f).

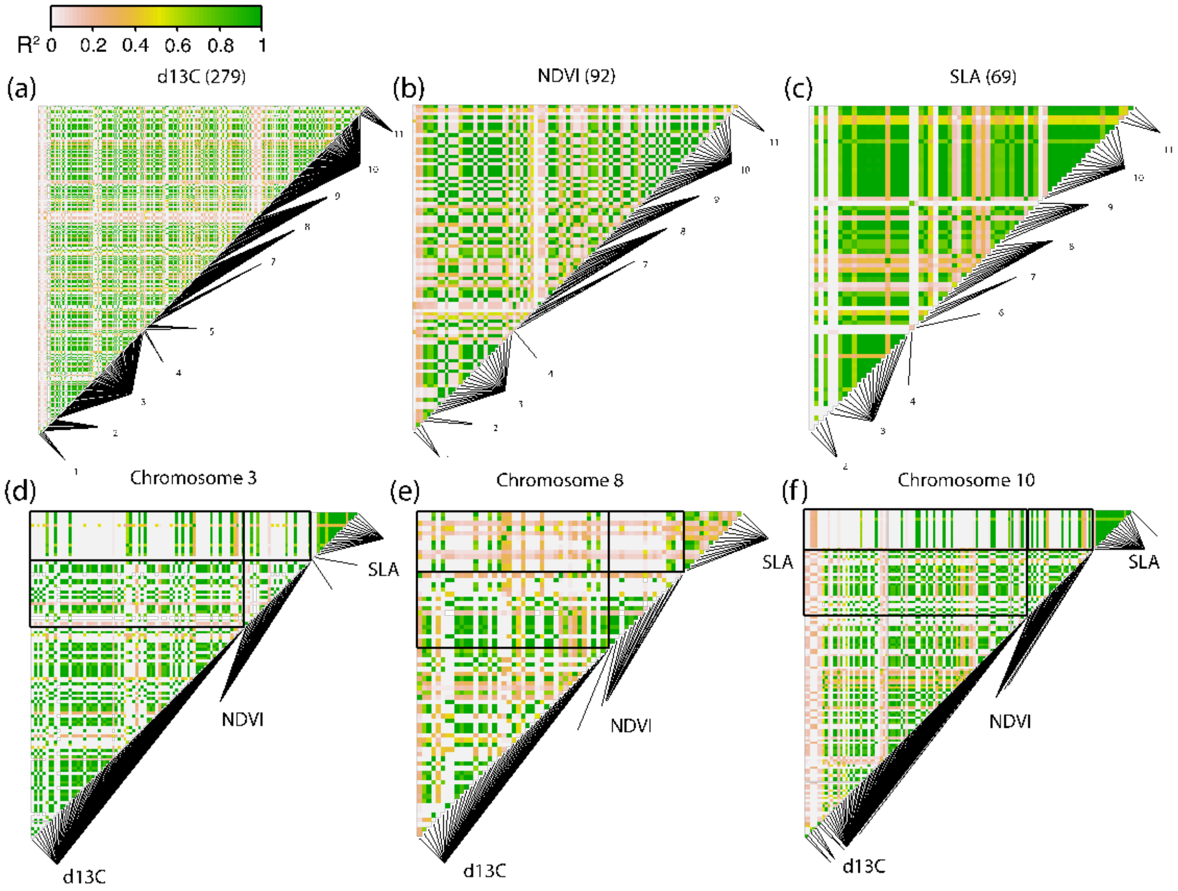

Figure S8. Full variant network output from the cape analysis. This view focuses on the structure of genetic interactions, regardless of variant position. Each node is one genetic SNP. Each is plotted as a pie chart with each trait as a specific piece of the pie. Significant effects are indicated by brown or blue section colouring, corresponding to positive (brown) or negative (blue) main effects with gray indicating no significant main effect. Interactions are shown as arrows between the nodes. Pie charts with both the brown and blue colours for different traits are indicative of antagonistic pleiotropy.

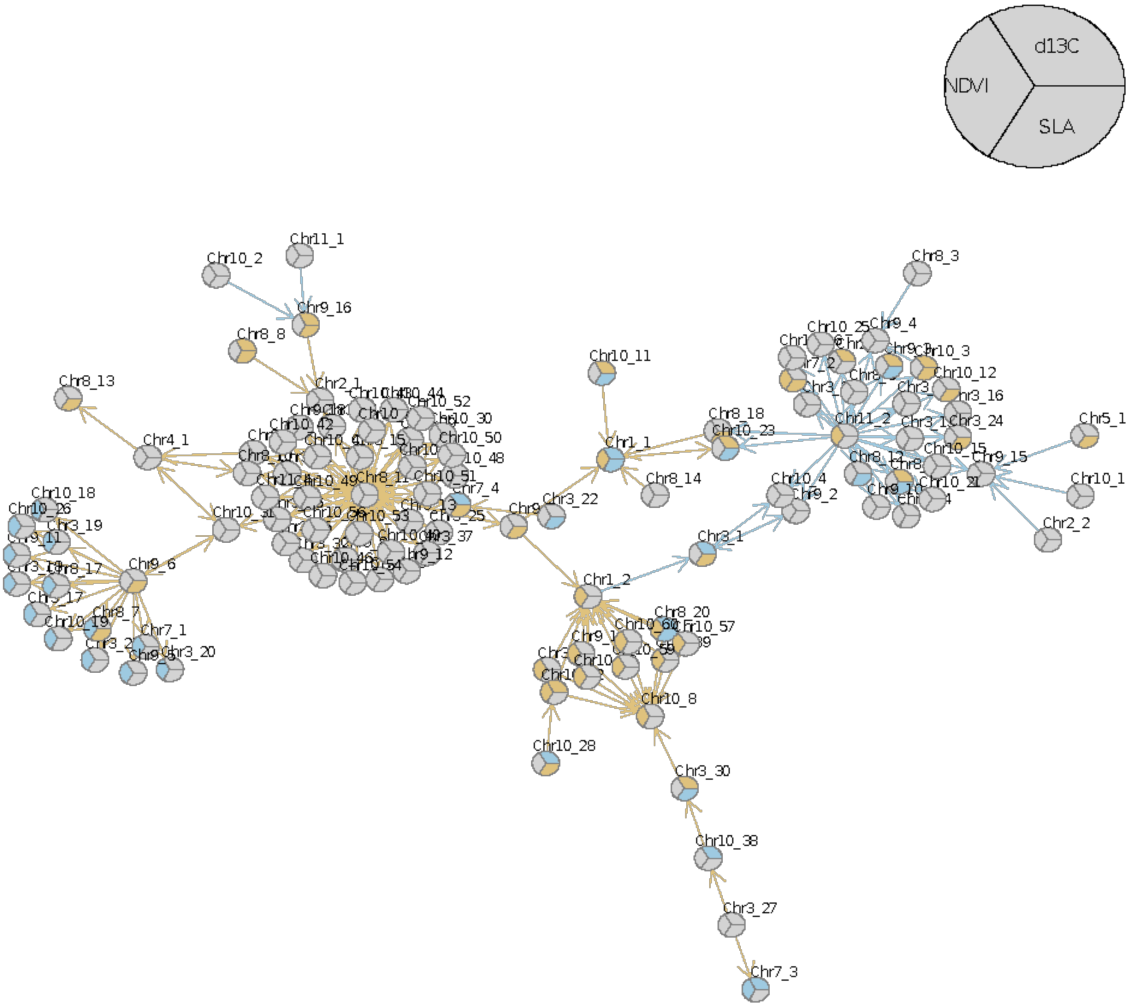

Figure S9. Gene networks from gene ontology enrichment analysis for all three categories for each trait.

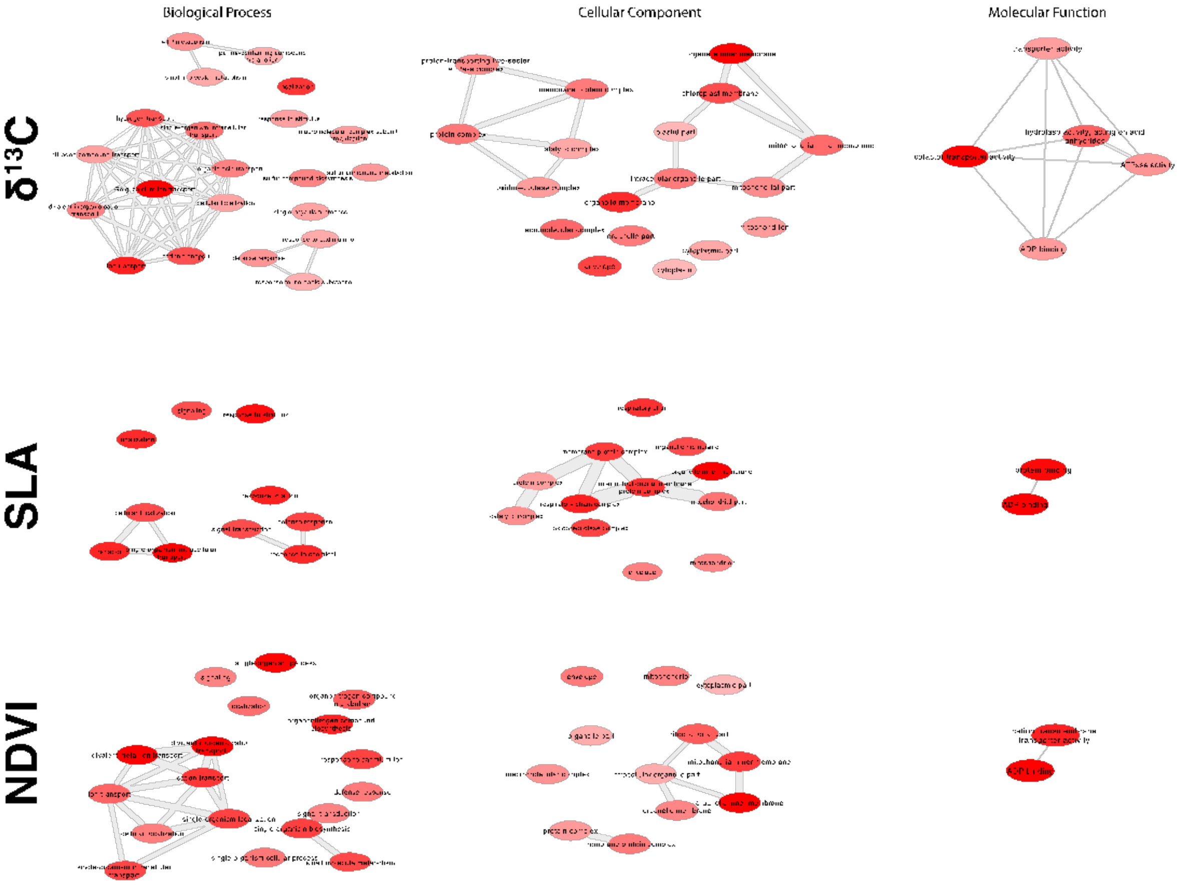

Figure S10 – Discriminant analysis for principal components for each trait.

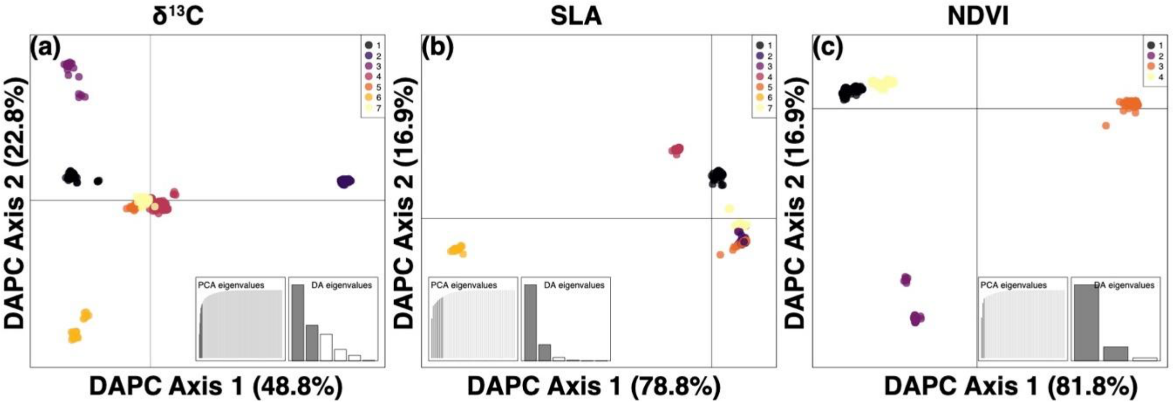

Figure S11 – Random forest analysis for Normalized Difference Vegetation Index (NDVI) for climate variables (top row position 1) and genetic variation from significant SNPs associated with NDVI (top row position 2). Predicted distributions are given for climate and genomic prediction. The difference between the predictions are shown. Along with the distribution of genetic variance.

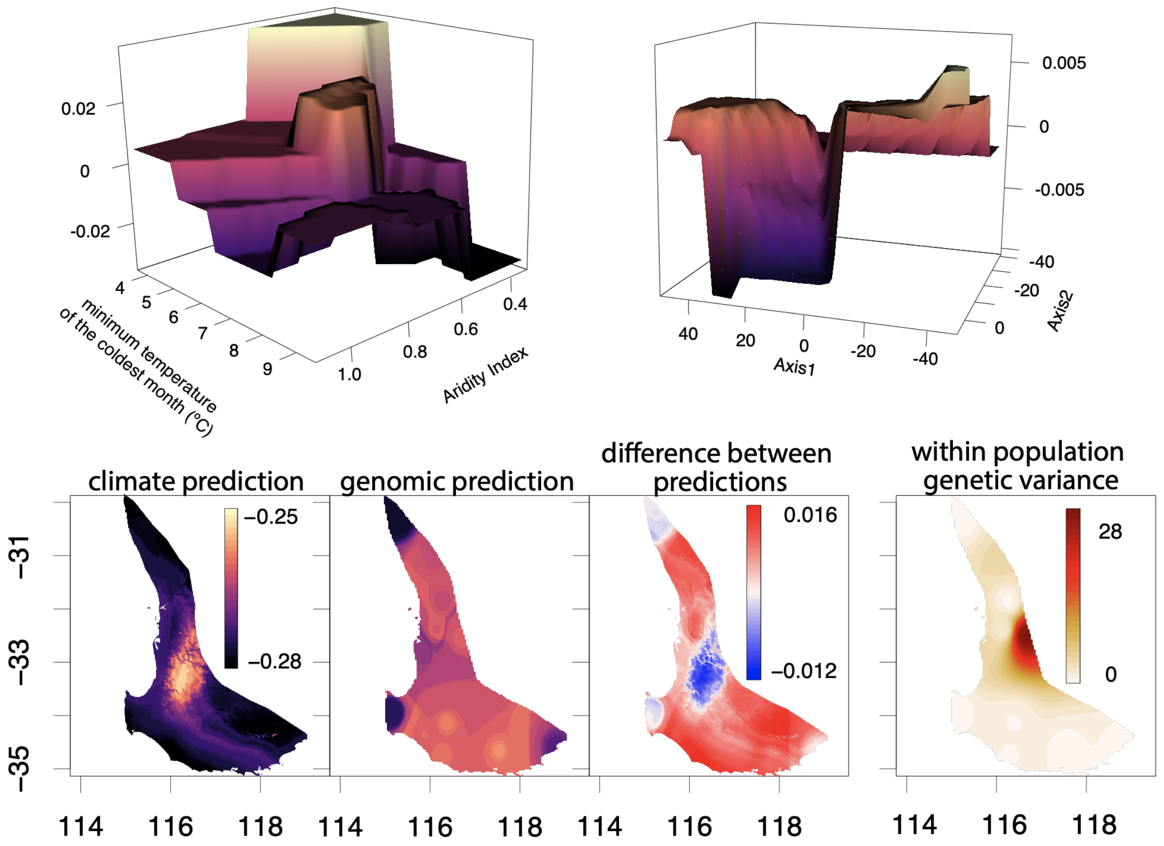

Figure S12 – Random forest analysis for specific leaf area (SLA) for climate variables (top row position 1) and genetic variation from significant SNPs associated with SLA (top row position 2). Predicted distributions are given for climate and genomic prediction. The difference between the predictions are shown. Along with the distribution of genetic variance.

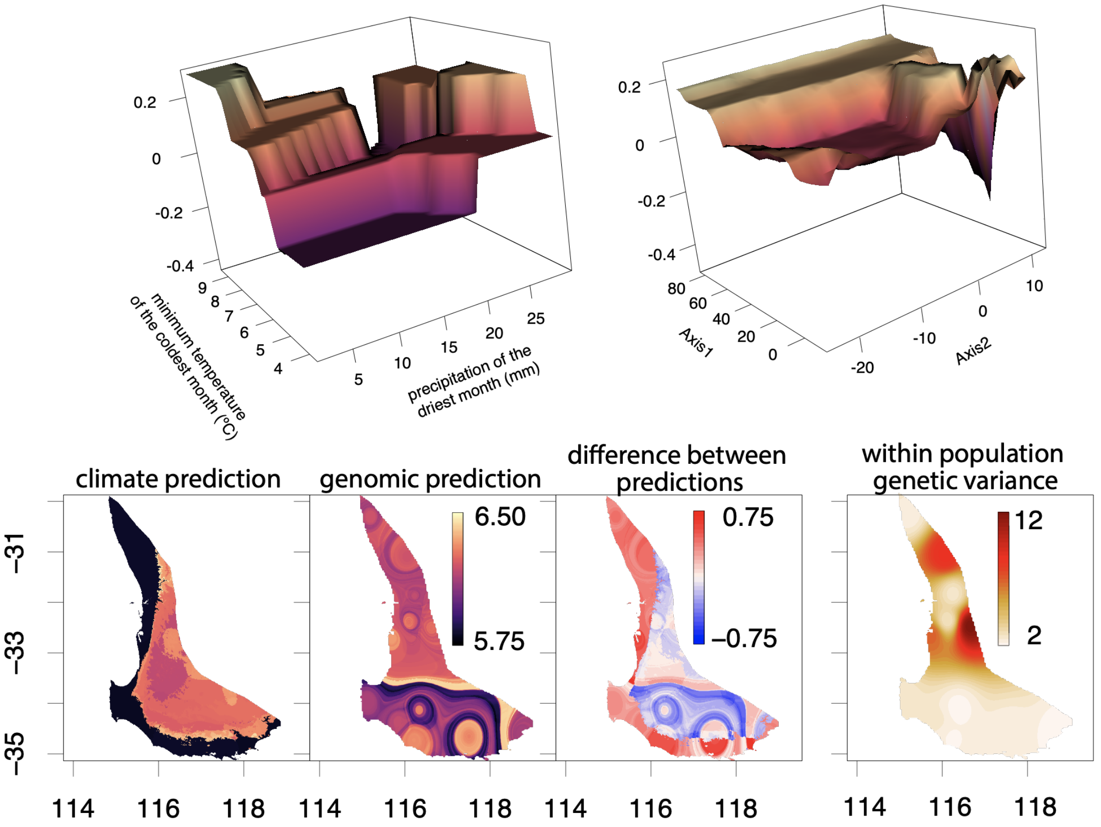

Figure S13. Manhattan plots showing the -log10(adjusted *P*-value) for precipitation of the driest month (BIO14).

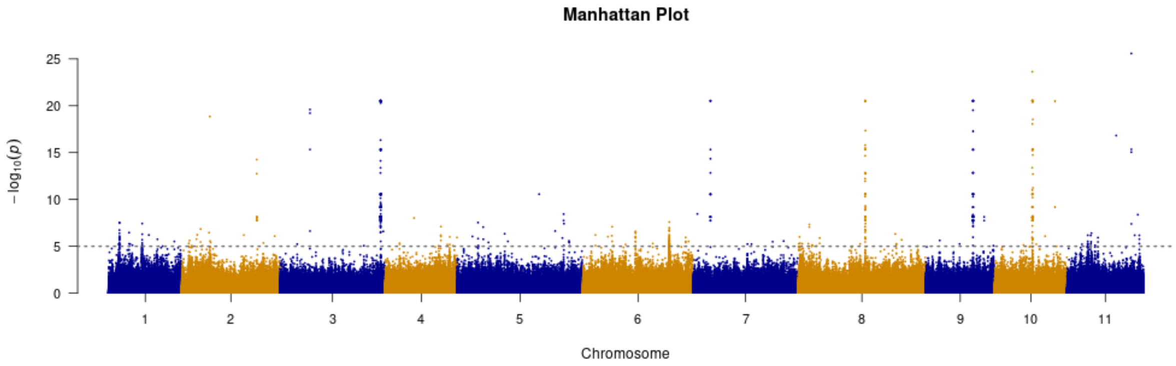

Figure S14. Location of genetic variation (estimated from standard deviation of the DAPC eigenvalues) associated with the SNPs significantly associated with precipitation of driest month (BIO14).

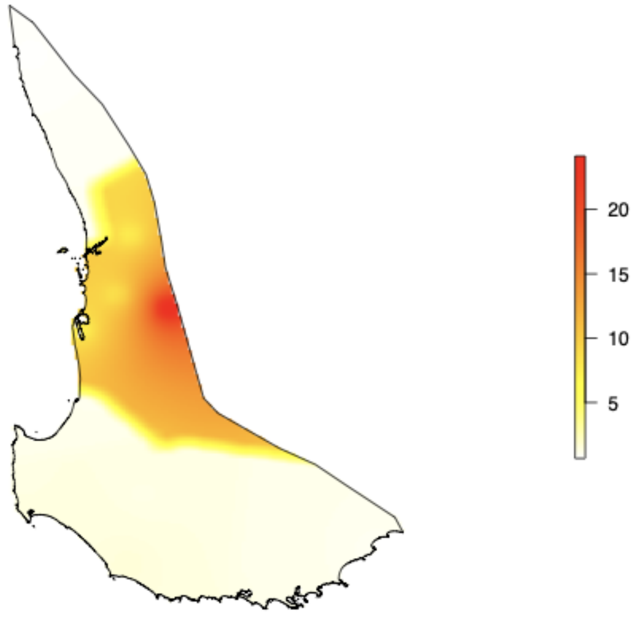

Figure S15 – Linear regressions between the number of SNPs (x-axis) and linkage disequilibrium (measured as halfmax) to if the number of SNPs per window was descriptive of measured linkage disequilibirum.

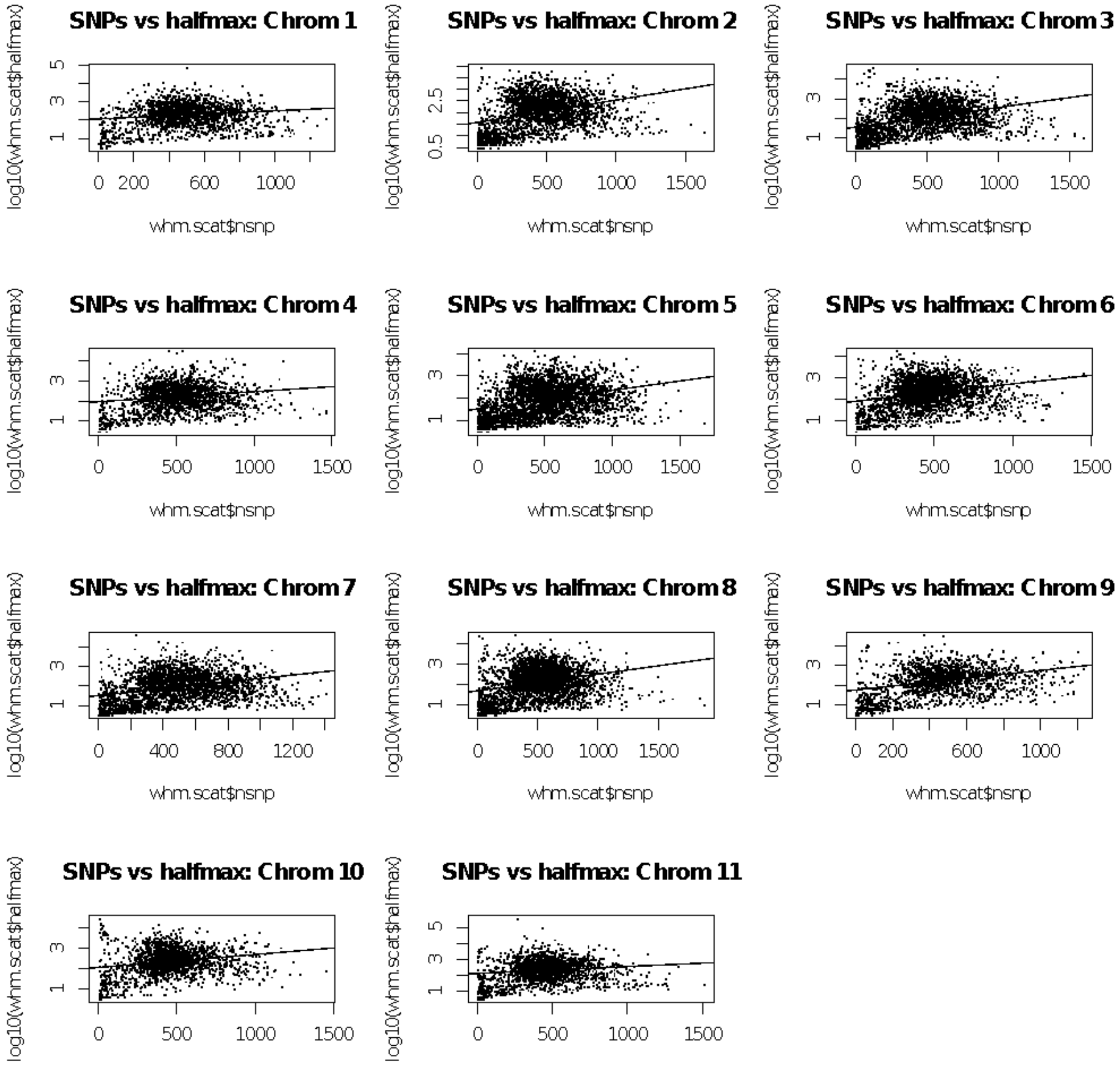

Figure S16 – Trait permutation to test if the GWAS associations are due to chance with $\delta^{13}C$. Manhattan plot 1 is the plot shown in Figure S8, manhattan plot 2 is permutation of trait values but keeping the family structure, and manhattan plot 3 is completely random permutation of trait values.

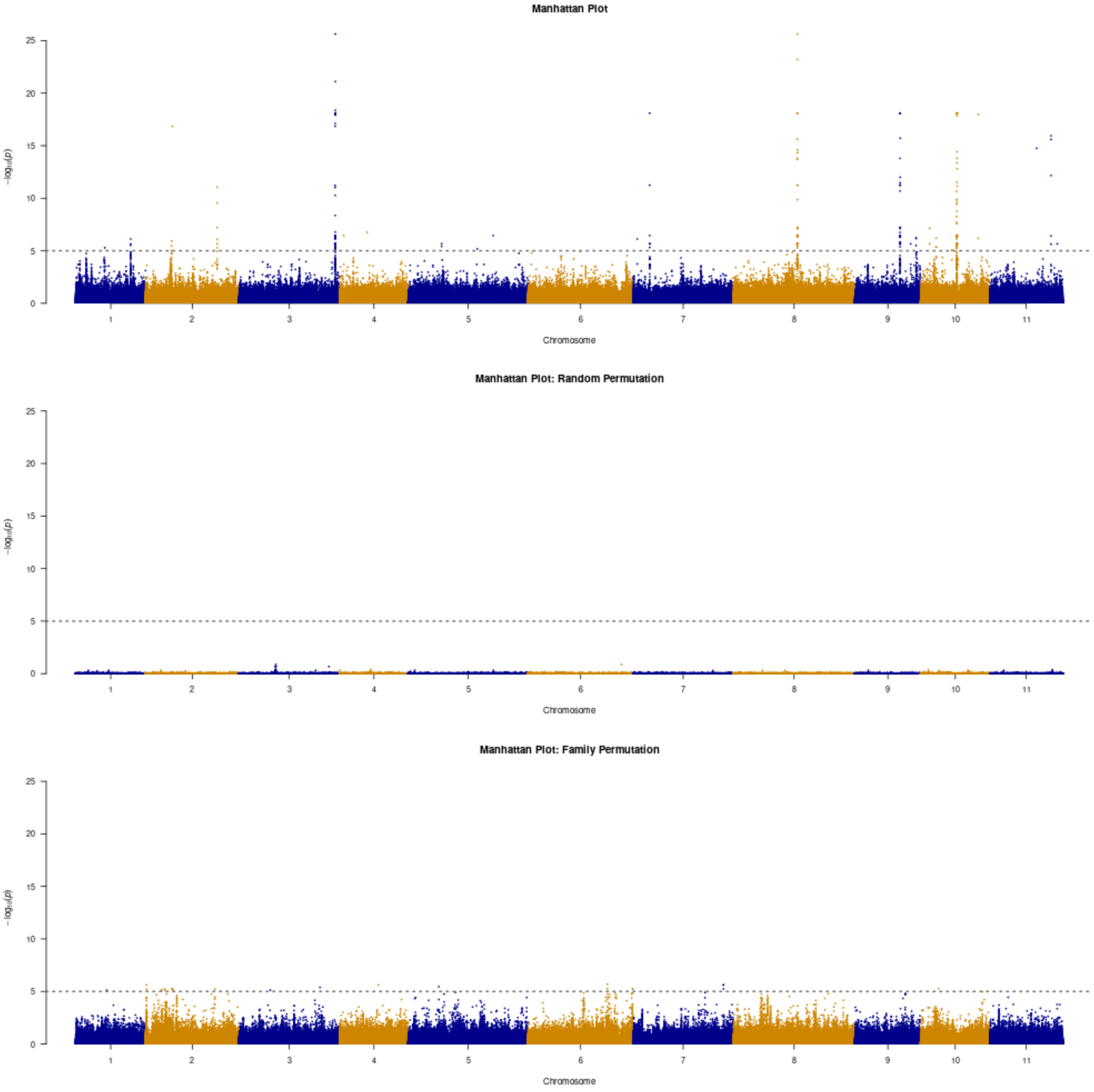

Figure S17 – Trait permutation to test if the GWAS associations are due to chance with NDVI. Manhattan plot 1 is the plot shown in Figure S8, manhattan plot 2 is permutation of trait values but keeping the family structure, and manhattan plot 3 is completely random permutation of trait values.

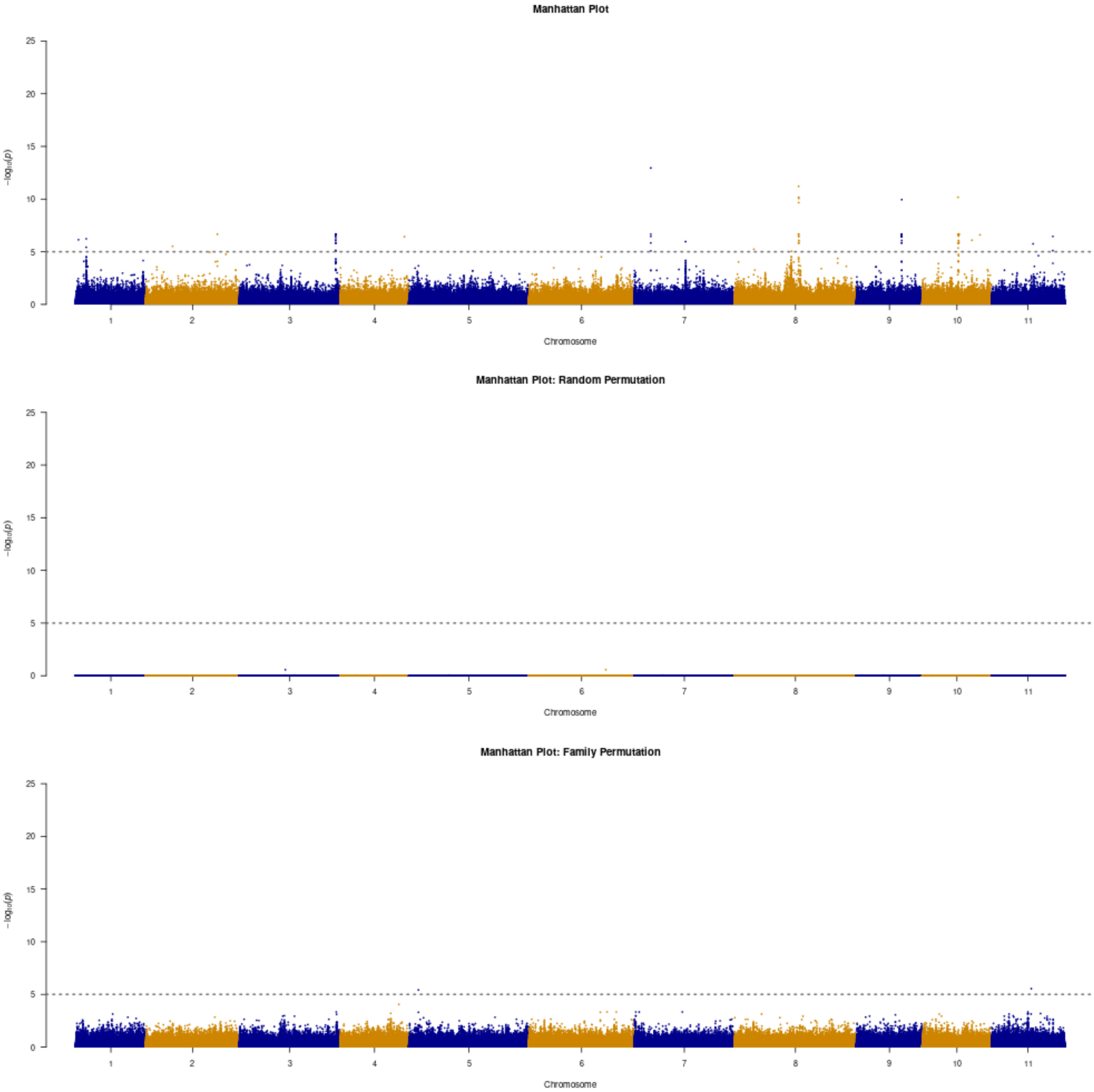

Figure S18 – Trait permutation to test if the GWAS associations are due to chance with SLA. Manhattan plot 1 is the plot shown in Figure S8, manhattan plot 2 is permutation of trait values but keeping the family structure, and manhattan plot 3 is completely random permutation of trait values.

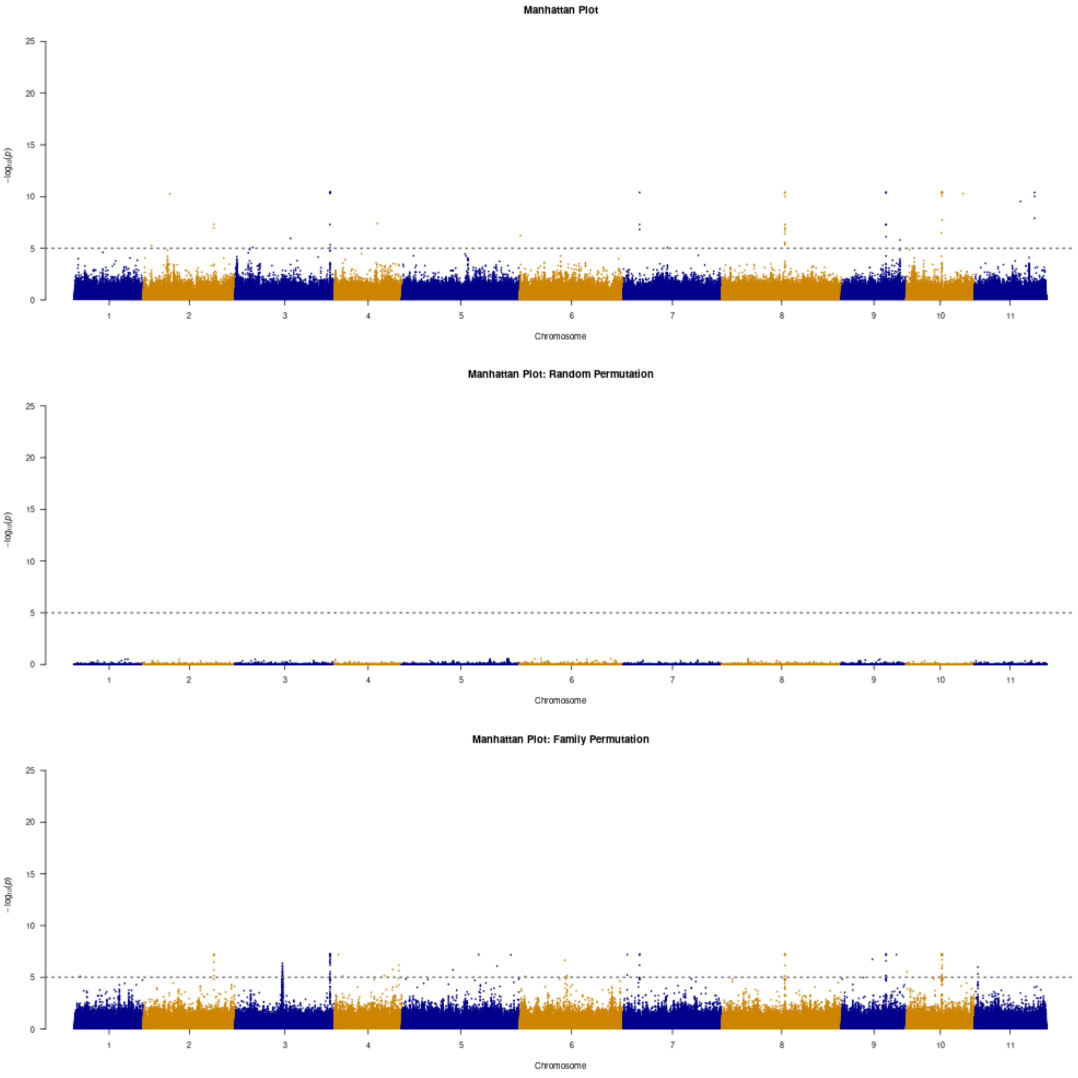

Figure S19 – Image plot of the kinship matrix.

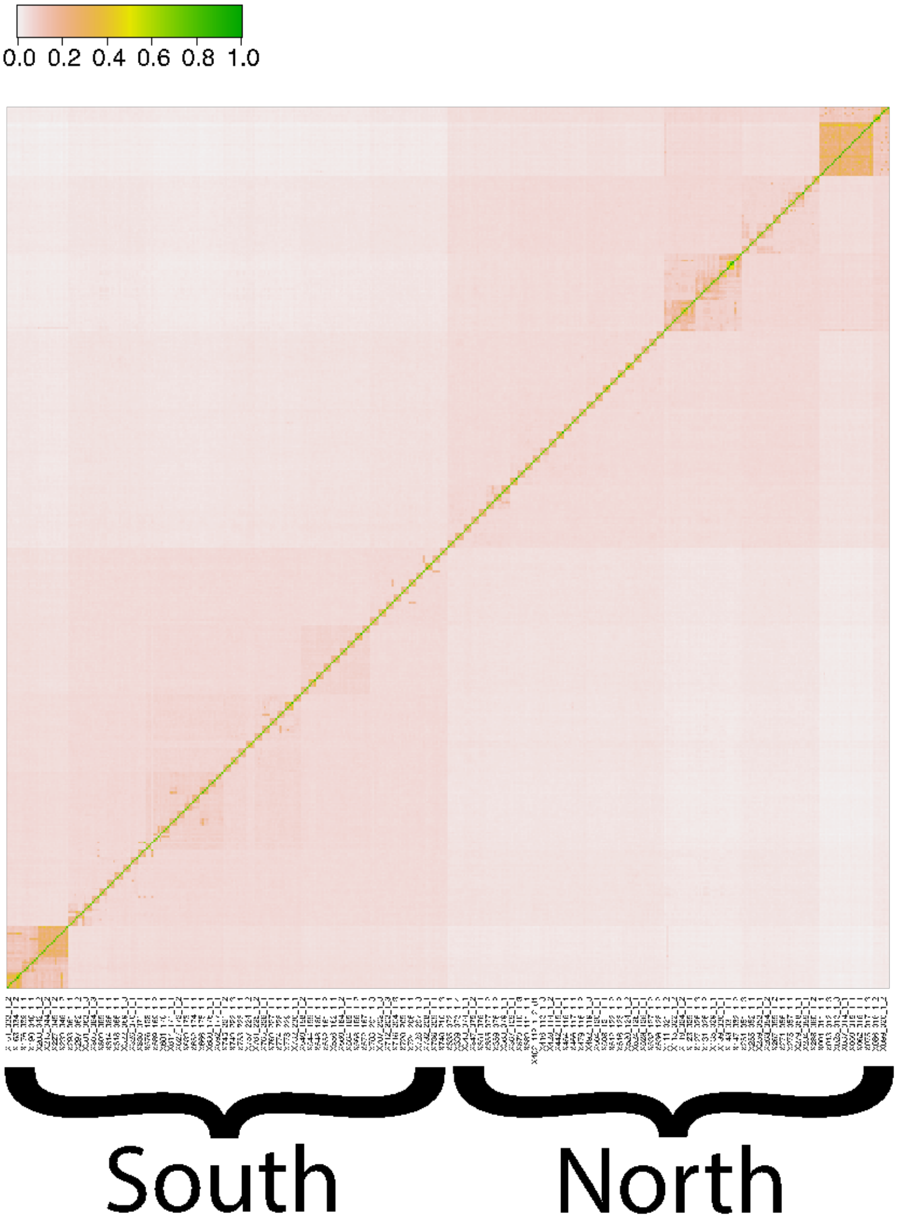

Figure S20 – Local genome population structure for the three important chromosomes calculated in 1000 SNP windows. Column two is coordinate 1 (y-axis) version window number (x-axis) and column three is coordinate 2 (y-axis) versus window number. Dashed grey line indicates the window in which peaks with significantly associated *P*-values occur.

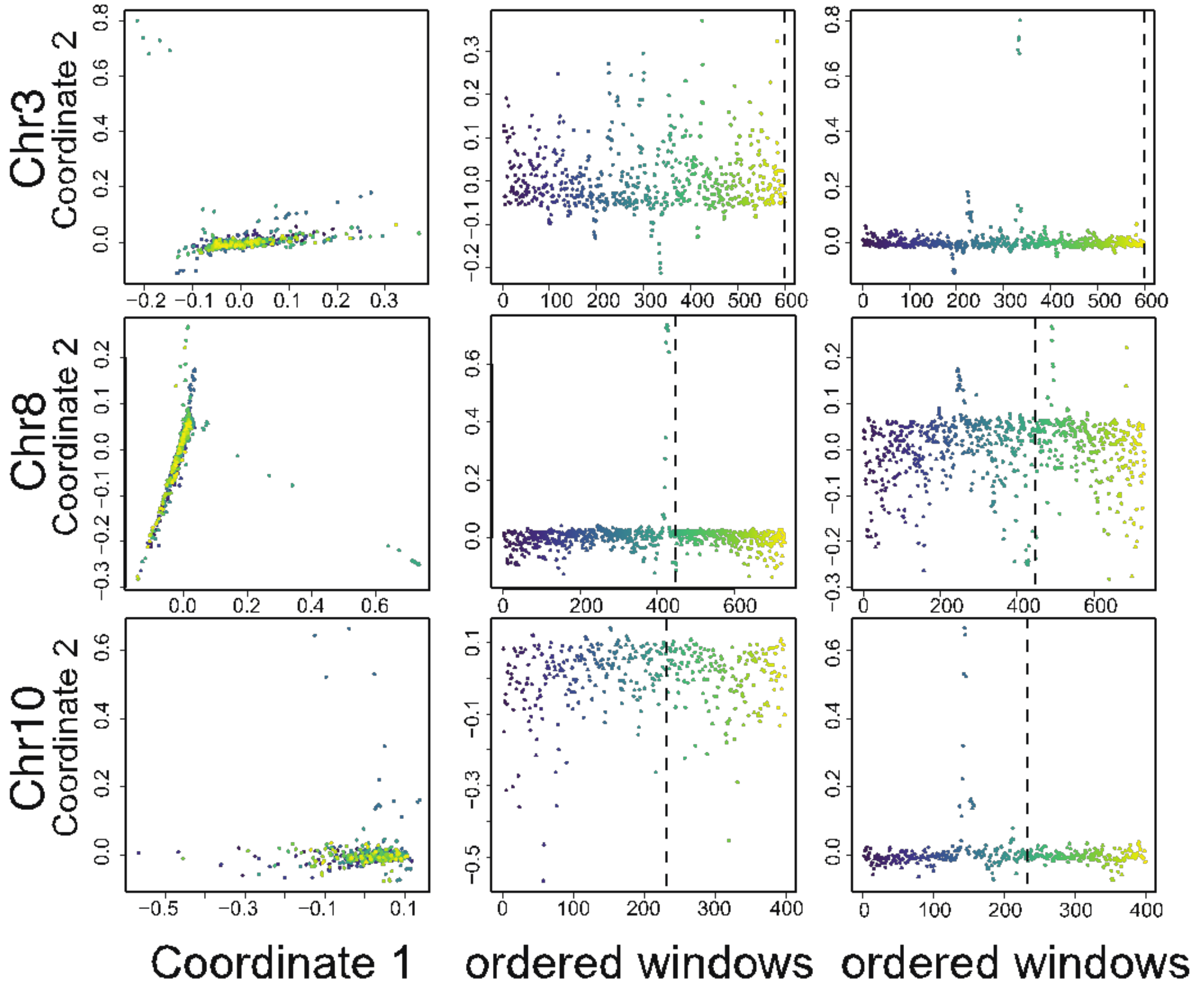
